# Evolutionary Trajectories of Ciprofloxacin Resistance in *P. aeruginosa* Lung Biofilms: Mutation Dynamics, Metabolomic Shifts, and Collateral Sensitivity

**DOI:** 10.64898/2026.05.07.723426

**Authors:** Doaa Higazy, Ke-Chuan Wang, Lene Bay, Steen Seier Poulsen, Pernille Rose Jensen, Claus Moser, Oana Ciofu

## Abstract

The evolution of antimicrobial resistance (AMR) in chronic biofilms is often viewed as a unidirectional path toward higher fitness, yet the metabolic constraints governing these trajectories remain poorly understood. We performed a four-passage evolution experiment using a murine lung biofilm model to assess the impact of prolonged ciprofloxacin (CIP) exposure on resistance and host response. This approach integrated population-level adaptive dynamics, whole-genome sequencing (WGS), and NMR-based metabolomics, alongside histopathology and cytokine analysis. Prolonged CIP treatment accelerated resistance, with isolates reaching MICs of 8–12 mg/L (a 32- to 48-fold increase) by the fourth passage. WGS revealed distinct evolutionary trajectories: control isolates accumulated metabolic and regulatory mutations without susceptibility changes, while CIP-treated isolates exhibited a stepwise progression from metabolic adaptation to high-level resistance, marked by early *nfxB* and late *gyrA* mutations. Metabolomic profiling revealed progressive divergence, with PCA identifying the *nfxB* genotype as the primary driver of variation (49.1% of variance). This resistant metabolic state was characterized by the depletion of central carbon metabolites, including glucose and tyrosine, alongside the accumulation of essential amino acids. Importantly, these changes were accompanied by a distinct trade-off; high-level CIP resistance triggered collateral sensitivity to tobramycin and aztreonam. While CIP treatment ultimately reduced neutrophilic inflammation (*p* = 0.011) and mucin production (*p* = 0.0496), early-passage lungs exhibited transient elevations in pro-inflammatory cytokines (CXCL2, MMP2, TNF-α). In conclusion, the adaptive trajectory to CIP resistance involves metabolic rewiring and collateral sensitivity, offering a framework to exploit the evolutionary costs of resistance in chronic biofilm infections.

## Introduction

The global escalation of antimicrobial resistance (AMR) is rapidly transforming manageable clinical conditions into untreatable crises. For patients living with chronic respiratory diseases, such as cystic fibrosis (CF) and bronchiectasis, this trend represents an existential threat as the pathogens they harbor evolve faster than our therapeutic interventions can keep pace (1). *Pseudomonas aeruginosa* stands at the center of this challenge; it is a remarkably versatile opportunistic pathogen that employs a complex array of intrinsic and acquired resistance mechanisms to persist within the hostile, nutrition-deprived environment of the lung and form biofilms (2). *P. aeruginosa* responds to the pressure caused by antimicrobial drugs by evolving and by its ability to survive and tolerate stress conditions. A research study assessed the tolerance and resistance to meropenem, ciprofloxacin (CIP), and tobramycin in *P. aeruginosa* isolates from chronic CF lung infections over 40 years of evolution (3).

Interestingly, tolerance was positively selected in the CF lung and may act as a precursor to resistance. However, the relationship between tolerance and resistance varied among patients and clone types, making it unpredictable (4). Therefore, studying the evolutionary trajectories and genomic diversity of AMR in *P. aeruginosa* is far from fully investigated and needs further attention to be able to prevent or circumvent the development (5). Still, the emergence of AMR is increasingly recognized not as an isolated genetic event, but as a complex crosstalk between signal transduction systems and metabolic networks. These regulatory loops allow pathogens like *P. aeruginosa* to sense nutritional shifts in the lung environment and dynamically adjust their metabolic flux to support AMR mechanisms (6). Consequently, characterizing the longitudinal transition from metabolic tolerance to stable genetic AMR is crucial for identifying potential vulnerabilities in the pathogen’s adaptive landscape. The survival of ESKAPE pathogens—*Enterococcus faecium*, *Staphylococcus aureus*, *Klebsiella pneumoniae*, *Acinetobacter baumannii*, *Pseudomonas aeruginosa*, and *Enterobacter* species—in the harsh, nutrition-deprived environment of the biofilm is facilitated by their remarkable metabolic versatility (7). When primary carbon sources are exhausted, these bacteria frequently undergo a metabolic switch—such as the activation of the glyoxylate shunt—to utilize alternative substrates for gluconeogenesis (8). While the glyoxylate cycle is a known focal point for AMR development due to its role in persistence, the broader relationship between central carbon flux and the evolution of high-level AMR remains poorly understood. Recent advances in metabolic state-driven approaches suggest that the AMR metabolome can be exploited to restore bacterial susceptibility (9). In this context, characterizing the longitudinal metabolic shifts during the evolution of *P. aeruginosa* is essential for identifying the metabolic trade-offs. To simulate the selective pressures of chronic respiratory infections, various experimental models have been developed to study *P. aeruginosa* adaptation. Specifically, *P. aeruginosa* embedded in alginate beads was shown to serve as a successful tool in driving chronic infections into the lungs of mice, which resembles the alginate-containing biofilms in a CF environment (10). Among the factors that affect the emergence of AMR is the dosage regimen, which includes dose level, dosing interval, and treatment duration (11, 12). Our previous study on CIP resistance evolution during four consecutive passages in an *in vivo* mouse model of chronic lung infection initiated with small bacteria embedded alginate beads showed that resistance was detected as early as the second passage, with mutations emerging in the *nfxB* and *mexZ* genes, which regulate efflux pumps. The study also highlighted the complex interplay between infection, antibiotic treatment, and the immune response (13). Building on our previous work (13), the present study sought to investigate the evolutionary consequences of prolonged CIP exposure in a chronic lung infection model. We utilized a modified dosing regimen of subinhibitory CIP—administered four times over 48 hours per passage—to better simulate the sustained selective pressures encountered during clinical therapy. Our primary aim was to map the longitudinal trajectory of *P. aeruginosa* as it transitions from initial tolerance to stable, high-level genetic resistance. We hypothesized that this rapid escalation in resistance is not merely a genomic event, but is governed by a metabolic canalization that imposes a high physiological cost on the pathogen. To test this, we integrated multi-passage genomic sequencing with non-targeted ^1^H-NMR metabolomic profiling and host histopathological analysis. By characterizing the interplay between *nfxB*-mediated efflux, central carbon reorganization, and host inflammatory responses, this study provides a framework for identifying evolutionary trade-offs and translational strategies that exploit the metabolic vulnerabilities of antibiotic-resistant biofilms.

## Materials and methods

### Bacterial strains, MICs, and animals

The reporter strain PAO1-*mCherry-PCD-gfp*+ (PAO1 background), which constitutively expresses red fluorescence via mCherry tagging and carries a chromosomal transcriptional fusion between the P*_mexCD_* promoter and *gfp,* was used. This setup allows green fluorescence to be triggered upon mutation of *nfxB*, a system previously developed in our lab and used in our previous study (14, 15). Female BALB/c mice (Janvier Labs, France), aged 10 – 11 weeks, were split into two experimental groups (control/placebo, untreated, n=40) and (CIP-treated, subcutaneous: s.c., n=40). Before the experiments began, the mice were acclimatized for 7 days at the Biotech Research & Innovation Centre at the University of Copenhagen. The animals were housed in a controlled environment with a 12-hour light-dark cycle, regulated temperature and humidity, and free access to food and water. Daily health observations were conducted, and disease progression was monitored regularly. To maintain consistency, the same researcher performed individual scoring for all mice.

### Alginate beads preparation

The alginate beads were prepared the same way as previously described (13). In summary, 1% alginate was prepared and then sterile-filtered. PAO1-*mCherry-PCD-gfp*+ strain was grown on an Luria-Bertani (LB) plate from a frozen stock, and a single colony was selected to start an overnight culture with an optical density of 2 at OD600. The bacterial culture of 20 ml was centrifuged, resuspended in fresh 5 ml LB medium, and mixed with the 1% alginate solution at a 1:20 ratio. Seaweed alginate beads were then prepared using the Nisco Var J30 Encapsulation Unit as previously described (16), utilizing a 0.250 mm nozzle with an alginate flow rate of 20 ml/h and a pressure setting of 35 mBar. Alginate bacterial mix (7 ml) was suspended in 100 ml of a gelling bath containing 0.1 M Tris-HCl and 12.5 mM CaCl2, stabilized for 1 hour during magnetic stirring, and then washed twice with 0.9% NaCl containing 0.1 M CaCl2. The colony-forming unit (CFU) counts were determined by plating the dissolved beads, and the stock was stored overnight at 4°C (Day 0).

### Lung infection and treatment with CIP (evolution cycle)

On day 1, embedded beads were adjusted to an inoculum of approximately 10^8^ CFU/ml beads in all the evolution passages. The mice were anesthetized subcutaneously (s.c.) with 0.1 ml/10 g mouse with a mixture of 1:1:2 (25%), a cocktail of hypnorm (fentanylcitrat 0.315 mg/ml and fluanisone 10 mg/ml) combined with (25%) midazolam (5 mg/ml), and (50%) sterile water (standard Hypnorm/Midazolam Mixture). Under anesthesia, the BALB/c mice were tracheotomized, and a volume of 40 μl of bacterial inoculum was delivered into the left lung with a curved-tipped needle. The incision was then sutured to allow healing (17), and the mice were maintained in a warm environment and closely observed until they regained consciousness. Subsequently, they received a subcutaneous injection of 1 ml saline to ensure rehydration and were administered analgesic buprenorphine (Temgesic 0.05-0.1 mg/kg s.c.) for pain management. On Day 2, the mice were scored and randomly split into two groups. One group was treated with CIP 0.25 mg (11.35 mg/kg) ×2 CIP subcutaneously with an 8-h interval for two consecutive days (a total of 4 administrations, Days 2 and 3), and the other control group received saline at the same time points. Rather than increasing peak concentrations, we extended the duration of exposure by administering four doses of CIP (11.35 mg/kg per dose) at 8-hour intervals over two consecutive days (Fig. S1&S2). On Day 4, the mice were euthanized, and the lungs were collected aseptically in Eppendorf tubes of saline 0.9% and homogenized using MagNaLyser and used for population analysis. The mouse lung infection model license number: 2024-15-0201-01742.

### Population analysis (cont. evolution cycle)

Luria–Bertani (LB) plates supplemented with CIP at different concentrations were prepared (0.1 mg/L to 8 mg/L). CFU counts for the whole population were determined for each sample and used to calculate the frequency of resistant subpopulations growing on CIP plates. We followed a selection criterion for the CIP-treated group, as after the first passage, the colonies that survived the highest CIP concentration were pooled (4 colonies), grown overnight, and used to form alginate beads to start the new passage, which was repeated till the fourth passage.

However, for the control (untreated group), the colonies were collected from CIP-free plates.

### Growth rate and lag phase extension

To evaluate growth kinetics, isolates selected from different evolutionary passages (C1=5, C2=7, C3=9, C4=5, T1=9, T2=10, T3=8, and T4=21) were tested for growth rates and lag phase extension. The control group colonies were randomly selected from CFU plates of homogenized lung samples. In contrast, colonies from the CIP-treated, evolved mouse lungs were obtained from CIP-supplemented plates used in the population analysis experiment. These colonies were chosen across a range of CIP concentrations, with selection based on distinct colony morphologies, aiming to capture the full diversity observed on both low- and high-CIP plates. Overnight cultures in LB were adjusted to an OD_600_ of 0.1 and diluted 1000-fold. Subsequently, 100 μl of each dilution was transferred in triplicate into a flat-bottom 96-well plate (Nunc, Thermo Fisher Scientific) and incubated in an Infinite F200 Pro plate reader (TECAN, Männedorf, Switzerland) at 37°C with shaking at 225 rpm for 24 h. Absorbance (OD_600_) was recorded every 20 minutes using Magellan V 7.2 software. The maximum specific growth rate (μ) was determined by calculating the slope of the linear regression of *ln*(OD_600_) over time during the exponential phase. Lag time (λ) was estimated by the intersection of the maximum slope line with the initial biomass concentration (OD_initial_), while the doubling time (*t_d_*) was calculated using the standard relationship 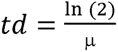.

### MIC determination

The minimum inhibitory concentration (MIC) for all the selected evolved isolates was determined. Overnight cultures were diluted to 10^−4,^ and 1 ml of the diluted culture was poured onto a blood agar plate, with the excess liquid discarded. The plates were allowed to dry, and the E-test (bioMérieux) strips were fixed in the middle of the plate. MICs for the background strain were 0.094 mg/L for ciprofloxacin and 2 mg/L for both tobramycin and aztreonam.

### Growth kinetics and time-kill analysis for tolerance testing

To characterize the response of evolved isolates to CIP, we performed both growth-based inhibition assays and time-kill kinetic studies. For growth inhibition, isolates from control and early-passage CIP-treated groups were cultured overnight in LB, adjusted to an OD of 0.1, and subcultured at 37°C until reaching mid-exponential phase (OD_600_ = 0.4). These cultures were then diluted 100-fold and transferred to 96-well microtiter plates containing a gradient of CIP concentrations. The plates were incubated in an Infinite F200 Pro (Tecan) at 37°C for 24 h to monitor optical density and survival across varying antibiotic pressures.

In parallel, time-kill assays were conducted to evaluate tolerance levels relative to the ancestral *P. aeruginosa* PAO1 strain. Standardized cultures were inoculated into 25 ml flasks containing 10 ml of LB and challenged with different concentrations of CIP (or an antibiotic-free control). Over a 24-hours (at 0, 0.5, 1, 2, 4, 6, and 24 h), 100 μl aliquots were collected, serially diluted, and 10 μl of each dilution was spotted onto LB agar plates for colony counting. Survival was quantified as the change in log_10_(CFU/ml) over time, allowing for the differentiation of killing kinetics and the identification of tolerant phenotypes among the evolved isolates.

### Collateral sensitivity

Evolved isolates were collected from various CIP plates with different concentrations for population analysis and from different passages. The isolates were passed twice in the LB medium in the absence of antibiotics. The MICs of ciprofloxacin, aztreonam, and tobramycin were determined for these isolates using E-test (bioMérieux SA, France). The study included 9 isolates from the first passage, 9 from the second passage, 7 from the third passage, and 18 from the fourth passage.

### Cytokine measurements

Cytokine levels in lung homogenate supernatants were quantified using a multiplex bead-based immunoassay (Mouse Magnetic Luminex Assays, R&D Systems, Abingdon, UK) according to the manufacturer’s instructions. Samples from infected and control mice were analyzed in technical duplicates. The analytes measured included a panel of core inflammatory mediators (CXCL2, CCL2, IFN-γ, TNF-α, G-CSF, GM-CSF, IL-1β, and IL-10). In addition, TWEAK (TNF-like weak inducer of apoptosis), S100A9, and MMP-2 were quantified to evaluate specific pathways of tissue injury and repair; specifically, S100A9 was included as a marker of alarmin-mediated innate activation, TWEAK for its role in modulating inflammation-induced cell death, and MMP-2 to assess extracellular matrix remodeling and structural integrity. Data acquisition was performed using a Luminex 200 platform (Luminex Corp., Austin, TX, USA), and cytokine concentrations were calculated from standard curves generated with recombinant standards.

### Histopathology

A total of four background control mice (infected with sterile alginate beads), six control mice from the fourth passage (infected with alginate beads embedded with bacteria and left untreated), and five infected mice treated with ciprofloxacin were euthanized. The trachea was canulated, and then 4% paraformaldehyde (PH 7.4) was injected into the lungs for 5 minutes and then excised and moved to a tube filled with paraformaldehyde and stored at 4°C overnight, and finally paraffin-embedded. The lung center was targeted by cutting tissue sections (4 μm) perpendicular to the surface. The lung sections were stained with Hematoxylin and eosin (H&E), as well as Periodic Acid-Schiff (PAS), dehydrated in 99% ethanol, and mounted (18). The PAS-stained slides were used to assess the presence of goblet cells, and mucus production in the Clara cells, while sections with H&E staining were used to assess the degree of inflammatory changes, including neutrophilic inflammation. The lung slides were assessed by a trained pathologist blinded to the grouping of the samples. A scale from 1 to 4 of the PAS-stained slides was employed using the following evaluation criteria: (1) a few Clara cells with a PAS-positive luminal edge, (2) PAS-positive edge, and quite a few Clara cells producing mucus in larger bronchioles, (3) PAS-positive edge and quite a few Clara cells in medium-sized bronchioles as well as a few goblet cells, and (4) several distinct goblet cells besides mucus containing Clara cells. A scale from 1 to 5 of the H&E stained slides was employed using the following evaluation criteria: (1) Neutrophils in a few alveoli, confined areas, (2) Neutrophils in alveoli in larger areas, (3) Neutrophils in all areas, (4) Neutrophils in all areas accompanied by condensation of the lumen, and (5) Changes corresponding to score 4 plus exudate in the bronchiole lumen.

### PNA-FISH staining

Paraffin-embedded mice lung tissue slices on the slides underwent deparaffinization through a standardized protocol (19), which included the following steps: (i) two washes in xylene for 5 minutes each, (ii) two washes in 99% ethanol for 3 minutes each, (iii) two washes in 96% ethanol for 3 minutes each, and (iv) three washes in Milli-Q water for 3 minutes each. The tissues were then incubated for 90 minutes at 55°C with a PNA-FISH-TexasRed (red) conjugated probe targeting a specific universal bacterial (UniBac) 16S ribosomal RNA (AdvanDx, Woburn, MA, USA). The slides were incubated in wash buffer (AdvanDx, Woburn, MA, USA) for 30 minutes at 55°C, followed by counterstaining with 4’,6-diamidino-2-phenylindole (DAPI) (Life Technologies, Oregon, USA) at a concentration of 3 μM for 15 minutes at room temperature. Finally, the slides were rinsed in Milli-Q water, covered with mounting media (Prolong Gold, Life Technologies, Oregon, USA), and sealed with a coverslip and clear nail polish (20).

### Microscopy

The slides prepared with PNA-FISH were imaged using a (Zeiss LSM 880, Carl Zeiss, Oberkochen, Germany) confocal laser scanning microscope. Fluorescence images were captured using excitation wavelengths of 405 nm and 594 nm, with emission wavelengths ranging from 410 to 483 nm (blue) and 599 to 690 nm (red). Images were visualized in 3D using Imaris 10 software (Bitplane AG, Zurich, Switzerland) and saved. Lung specimens were examined using a Zeiss Axio Imager A1 microscope equipped with an AxioCam ICc3 digital camera. Images were captured and processed using Zeiss ZEN 2 (blue edition) software.

### Whole genome sequencing-WGS

Bacterial isolates from the four control passages (C1–C4) and ciprofloxacin-treated passages (T1–T4) were selected (C1=4, C2=5, C3=4, C4=5, T1=4, T2=5, T3=5, and T4=7 from different populations (C1=4, C2=5, C3=4, C4=5, T1=4, T2=5, T3=5, and T4=7 and whole-genome sequenced. Selection criteria were established to account for biological variability among individual mouse lungs and for phenotypic diversity in MIC levels across isolates in distinct evolutionary passages. An overnight culture was prepared from each isolate, and DNA was extracted using a DNeasy Blood & Tissue kit (Qiagen, Venlo, Netherlands). DNA sequencing was obtained by Illumina technology (Eurofins Genomics, Ebersberg, Germany). The raw fastq reads were checked for quality using FastQC, and further aligned with the PAO1 reference genome (GenBank accession: AE004091) with bwa-mem and SAMtools. The variants were further called with Bowtie2 and BCFtools and annotated with SnpEff.

### ^1^H-NMR-based metabolomics and association analysis

#### a. Exometabolome profiling

To determine how genomic modifications translated into functional phenotypes, we performed non-targeted H-NMR metabolomics on cell-free supernatants from 37 selected *P. aeruginosa* isolates (C1=5, C2=7, C3=5, C4=6, T1=5, T2=4, T3=4, T4=7). This metabolic approach characterizes specific patterns of metabolite utilization and excretion by measuring the flux of small-molecule concentrations in the growth medium. This method provides a universal, unbiased profile of all metabolites above the detection threshold with high analytical precision, making it an ideal tool for identifying physiological shifts associated with antibiotic adaptation.

#### b. Sample preparation and acquisition

Metabolic activity tests were performed on the isolates used for WGS to construct an association network. All the isolates were cultured in LB broth at 37°C with shaking overnight. Following incubation, cultures were centrifuged, and the supernatants were filtered through 0.2 µm filters to obtain cell-free supernatants. For NMR analysis, 500 µL of each cell-free supernatant was mixed with 100 µL of phosphate buffer (pH 7.4) containing 600 mM NaH□PO□, 2 mM maleic acid, and 0.1 mg/ml sodium 3-(trimethylsilyl)-1- propanesulfonate (DSS) in 60/40% D□O/H_2_O. One-dimensional ^1^H-NMR spectra were acquired at 298 K on a 500 MHz Bruker spectrometer (Bruker, Fällanden, Switzerland) using a standard pulse sequence (noesygppr1d) with the following parameters: TD 32768, 4 s interscan delay (d1), 1.7 s acquisition time, and 256 scans. All spectra were processed and analyzed using MestReNova software (version 15.0.1, Mestrelab Research, Santiago de Compostela, Spain). PCA analysis and unsupervised hierarchical clustering were performed with MATLAB R2025.

#### c. Genotype-phenotype association analysis

Metabolomic profiles and genomic mutation data were integrated into a unified association analysis for 37 isolates. Metabolite abundances were standardized using Z-score normalization and log-transformation (*log*1*p*) before analysis. To identify non-linear dependencies between regulatory drivers (e.g., *nfxB*) and metabolic outputs, we utilized Spearman’s rank-order association (*ρ*). For the comprehensive association matrix (Fig. S4) and targeted amino acid analysis (Fig. S5), data were visualized using the pheatmap R package. A targeted association network (Fig. 6b) was constructed using a force-directed layout, where edge weights represent the magnitude of the association (*ρ*).

### Statistical analysis

Data analysis utilized unpaired t-tests via Prism 10 (GraphPad Software, San Diego, USA). Lognormal Welch’s t-test was used for analyzing CFU counts. A multiple unpaired t-test was used for population analysis. Computational analyses were performed using R (version 4.5.2).

The principal component analysis (PCA) for the association analysis was conducted using the prcomp function, with 95% confidence ellipses overlaid to quantify genotype-specific separation. The significance of gene-metabolite associations was assessed using Spearman’s rank correlation coefficients (ρ). Hierarchical clustering for all heatmaps was performed using the Ward.D2 method on Euclidean distances to identify functional metabolic modules. The cytokines’ statistical analyses were performed using Prism. Values below the lower limit of quantification (LLOQ) were assigned the standard curve value of LLOQ, and values above the upper limit of quantification (ULOQ) were assigned the ULOQ. Two-group comparisons were performed using the non-parametric Mann–Whitney test.

## Results

### The Population Shifts and Resistance Emergence in Biofilm-Infected Lungs

To evaluate the emergence of resistance under sustained selective pressure, we utilized a four-passage murine lung infection model. We administered an intensified dosing regimen of four-CIP administrations (11.35 mg/kg) over two days with 8-hour intervals within each 24h period (Fig. S1 and S2). Quantitative bacteriology demonstrated shifting dynamics in bacterial burden across the sequential passages (Fig. 1a). In the first passage, CIP-treated mice exhibited higher CFU counts in their lungs compared to the untreated controls (*p*=0.0029). However, this difference was abolished in the second and third passages as the populations adapted. A significant divergence in lung bacterial load re-emerged in the fourth passage, with the CIP-treated group exhibiting lower CFU counts than the control (*p*=0.0055).

**Fig. 1:**
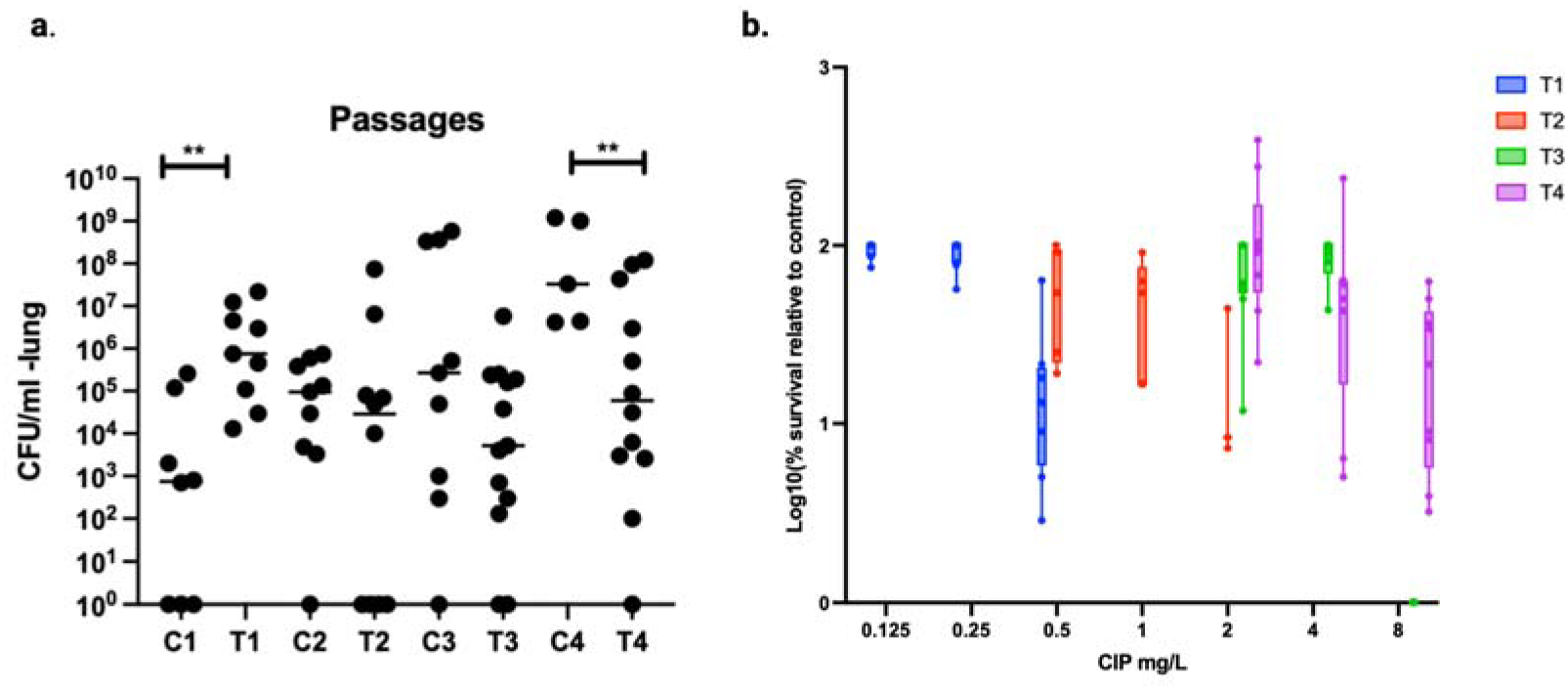
Population analysis profiles. **a.** Bacterial load in the lungs (CFU/ml) from individual lung homogenates across four passages in control (C1–C4) and ciprofloxacin-treated (T1–T4) groups. Statistical comparisons were performed using a log-normal–transformed Welch’s *t*-test (*p* < 0.05). **b.** Population analysis profile (PAP) of bacterial populations recovered from individual lungs across successive passages. The highest CIP concentration at which bacterial subpopulations survived increased across passages: 0.5 mg/L in the first passage, 2 mg/L in the second passage, and 8 mg/L in the third and fourth passages. Statistical analysis was performed using an unpaired *t*-test.

Phenotypic profiling of the recovered bacterial populations across different passages from mouse lung homogenates revealed a progressive escalation in CIP resistance, where the surviving subpopulation in the first passage initially exhibited survival at concentrations up to 0.5 mg/L (Fig. 1b). This threshold of phenotypic resistance increased steadily through the second and third passages until the fourth passage, where a significant (*p*= 0,001) fraction of the population survived at 8 mg/L CIP, representing a 16-fold increase in tolerated concentration compared to the initial survivors of the first passage.

### Isolate-Level Characterization of Resistance and Fitness Trade-offs

To confirm these population-level shifts, we determined the Minimum Inhibitory Concentrations (MICs) of representative isolates (Fig. 2a). A significant escalation in CIP resistance occurred between the first and second passages (*p* = 0.0153), with some isolates showing MICs of 4 and 8 mg/L in the second passage. MIC values increased further in subsequent passages, reaching a maximum of 12 mg/L by the fourth passage. However, no statistically significant difference was detected between the third and fourth passages, indicating that MIC levels plateaued at 12 mg/L (Fig. 2a).

**Fig. 2:**
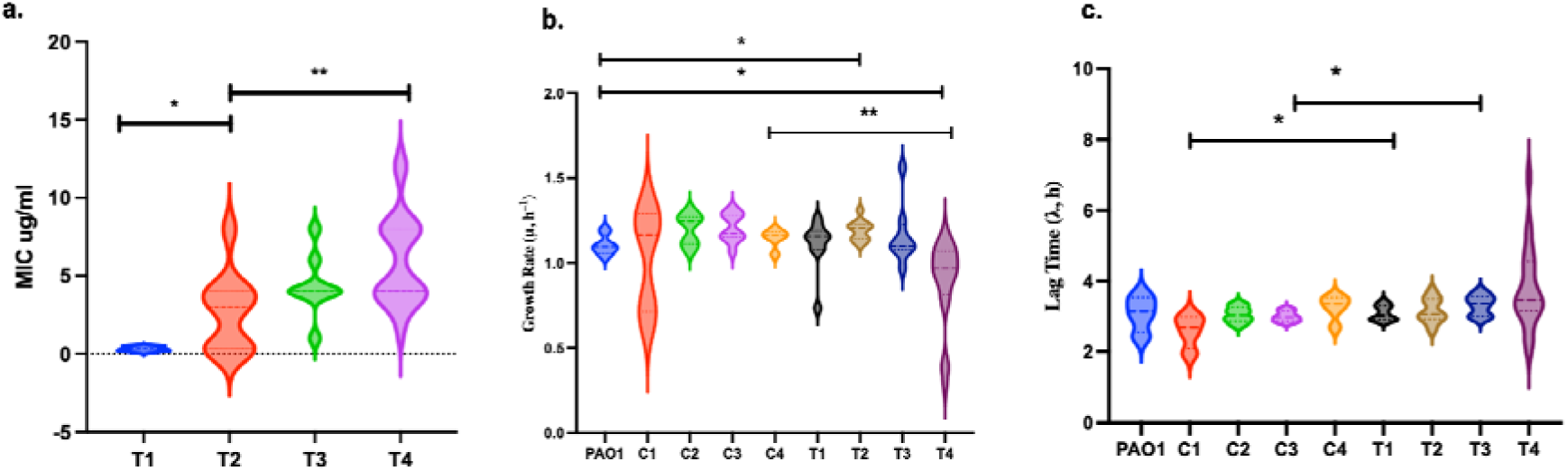
a. Minimal inhibitory concentrations (MICs) of CIP for bacterial isolates survived during PAP on LB plates supplemented with different concentrations of CIP in different passages (T1-T4). determined by E-tests. Violin plots illustrate the distribution and density of MIC values within the sampled population. b. The maximum growth rate (μ_max_) was calculated for the ancestral PAO1 and for isolates from control (C1-C4) and treated (T1-T4) passages. **c.** The extension of the lag phase (λ) was determined for all control (C1-C4) and treated (T1-T4) passages. All statistics were done with the Mann-Whitney U test, \**p* < 0.05, ** *p* < 0.01.

The stabilization of resistance level coincided with measurable shifts in bacterial fitness, characterized by altered planktonic growth kinetics in LB medium (Fig. 2b,c). Analysis of the maximum specific growth rate (*μ_max_*) revealed that while early evolved isolates (T1-T3) grew comparably to the ancestral PAO1, T4 isolates exhibited a significant reduction in *μ_max_* (*p* = 0.0184 vs. PAO1; *p <* 0.005 vs. C4 and T1–T3). This kinetic impairment was further characterized by a significant extension of the lag phase duration (λ) in T4 isolates compared to earlier passages, as in C3, T1 (e.g., vs. C3, *p* = 0.0081; vs. T1, *p* = 0.0328) (Fig. 2c).

To further dissect the survival mechanisms beyond static MIC measurements, we performed time-kill kinetics and growth profiles on isolates recovered from early passages (Fig. S3 and S4) and with low MIC levels (MIC < 0.5 mg/L). While the population-level data indicated an initial shift in resistance during the first passage (Fig. 1b), as observed in the time-kill analysis of the isolate (e.g., M5P1T05) (Fig. S3b), several isolates exhibit a tolerance-like phenotype (e.g., M2P2T05 and M3P2T1) as they were able to survive low CIP concentrations (0.1 and 0.2 mg/L) compared to the ancestral PAO1 (Fig. S3). Interestingly, isolates from the control passages (C1–C4) maintained killing kinetics and growth profiles close to PAO1 with a slight minimal difference in some isolates, confirming that the observed tolerance and subsequent resistance were driven strictly by the ciprofloxacin selection pressure rather than host-environment adaptation alone (Fig. S4).

### Evolutionarily developed adaptive mutations

To distinguish antibiotic-driven evolution from host-driven adaptation, we performed whole-genome sequencing (WGS) on isolates evolved in the absence of ciprofloxacin (CIP) across four sequential passages. These control isolates maintained stable MICs (≤0.2 µg/ml) but exhibited a progressive accumulation of mutations linked to metabolic remodeling and signal transduction (Fig. 3). Notably, the majority of the identified genetic modifications were concentrated within biosynthesis and metabolism, suggesting that the primary adaptive pressure of the murine lung involves significant nutrient and energy reorganization, also suggesting metabolic adaptations to *in vivo* environmental stresses, such as inflammation. Within this metabolic category, we identified mutations in genes governing polyamine catabolism (*pauA7*), amino acid biosynthesis (*argJ, hisF1*), and energy production (*pntB*, *pcaB*). Specifically, mutations in *pauA7* emerged by the first passage and achieved near-fixation across the sampled population by passage four. The adaptive landscape was further characterized by modifications to signal transduction pathways, most notably the mutations in the two-component regulator *parS* starting in passage three. Additionally, we observed mutations in transcriptional regulators (*gcdR*, PA1283), and cell wall/membrane components (*tolB*, mucB). *tolB* maintains outer membrane stability, while *mucB* regulates the alginate production and is crucial for mucoid biofilm formation. Despite the absence of drug pressure, a single isolate in the fourth passage acquired a mutation in the *mexY* efflux pump regulator, though this did not alter the CIP sensitivity profile (Fig. 3).

**Fig. 3:**
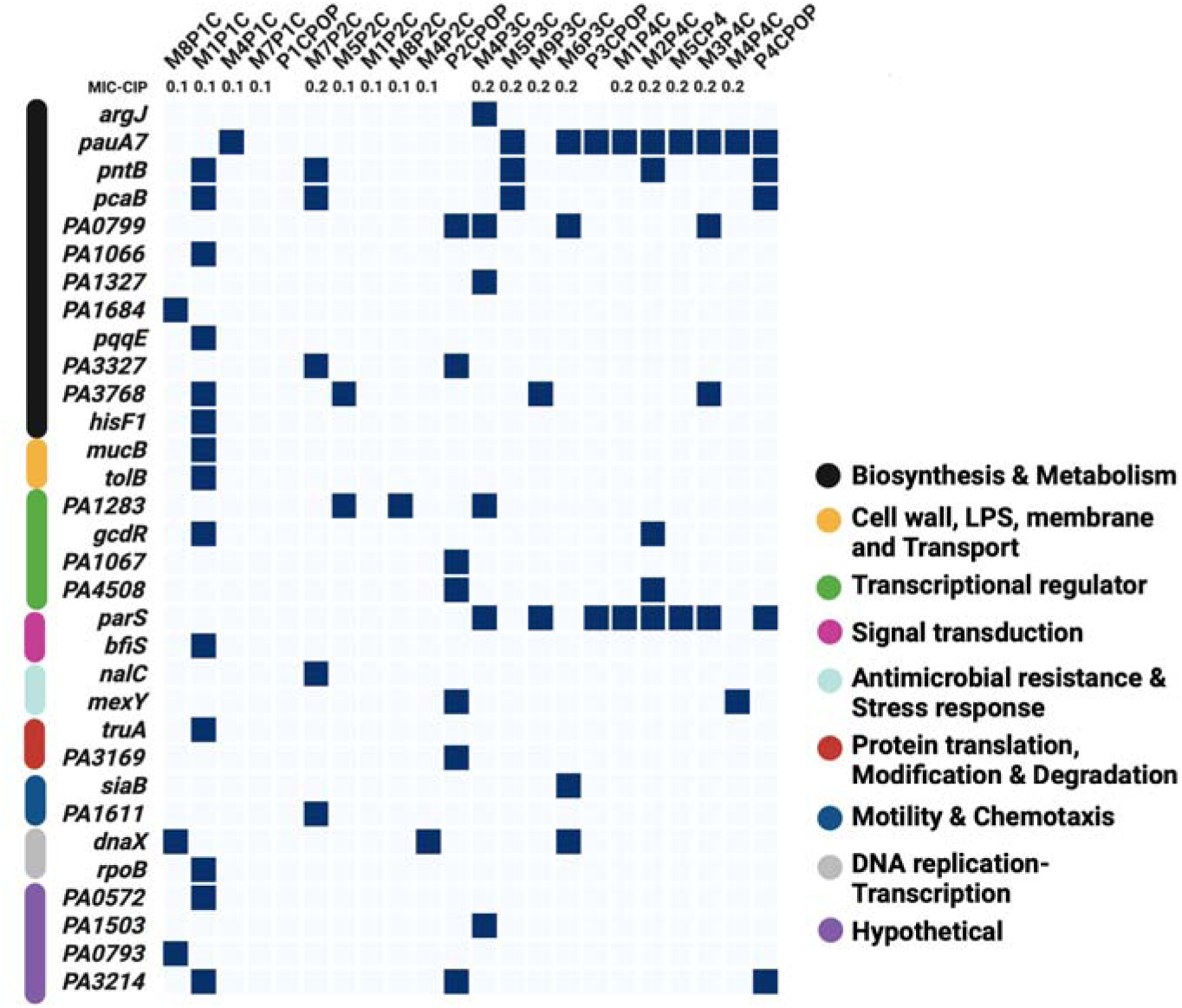
Genomic adaptation of *P. aeruginosa (*PAO1-*mCherry*-P*_CD_*-*gfp*+) to the murine lung in the absence of antibiotic pressure. Heatmap of mutational profiles in control isolates recovered across four sequential passages. Individual isolates are identified by mouse ID (M), passage number (P), and control group designation (C), with corresponding ciprofloxacin (CIP) MIC values (mg/L) indicated above each column. Blue cells denote the presence of nonsynonymous mutations within specific genes, categorized by functional groups as indicated by the vertical color bars.

Conversely, bacterial isolates recovered from mice treated with CIP exhibited a distinct evolutionary trajectory characterized by an initial phase of metabolic tolerance and structural adaptation, followed by resistance development. Across all passages, the majority of mutations were categorized in biosynthesis and metabolism, Cell wall, lipopolysaccharides (LPS), membrane, transport, and transcriptional regulators, mirroring the adaptive categories seen in the control passages but with treatment-specific targets. In early passages, mutations were in genes mainly related to metabolic and regulatory mechanisms, such as *pntB* (energy metabolism)*, pchE* (iron acquisition), and *aer2* (which play a role in chemotaxis). In addition to the mutations observed in *dnaX,* involved in the DNA replication process, and *folD,* involved in the folate pathway.

A critical shift toward high-level resistance occurred starting in the second passage, where the negative regulator of the MexCD-OprJ efflux pump *nfxB* was mutated in three out of five sequenced isolates, leading to increased drug efflux. In the third passage, the number of mutations in genes involved in antibiotic resistance increased. Such as mutations observed in *mexY* efflux and *nfxB* that were observed in all tested isolates. Mutations in genes involved in a number of metabolic pathways, including *pcaB,* were also observed, as well as mutations in genes involved in transcription regulation, such as *gcdR*. However, isolates from the fourth passage displayed the highest level of resistance development on the MIC level, correlating to the largest number of mutations that emerged in the different isolates, which shows that bacteria were highly adapted to the high concentrations of CIP by accumulation of mutations. By the fourth passage T4 of the treated group, this was complemented by the emergence of mutations in the primary target gene *gyrA,* which affect the fluoroquinolones’ binding efficacy, exclusively in isolates reaching the maximum MIC of 12 µg/ml (Fig. 4). This late-stage adaptation was further defined by a mutations in genes governing membrane integrity and envelope stress response (*tolB*, *mucB*), as well as translation and metabolic transcriptional regulation (*lepA*, *gcdR*). Mutations in genes like *tolB* play a role in membrane stability, and *lepA,* as a ribosomal factor, could all explain the adaptation gained by bacteria to survive CIP-imposed stress conditions.

**Fig. 4:**
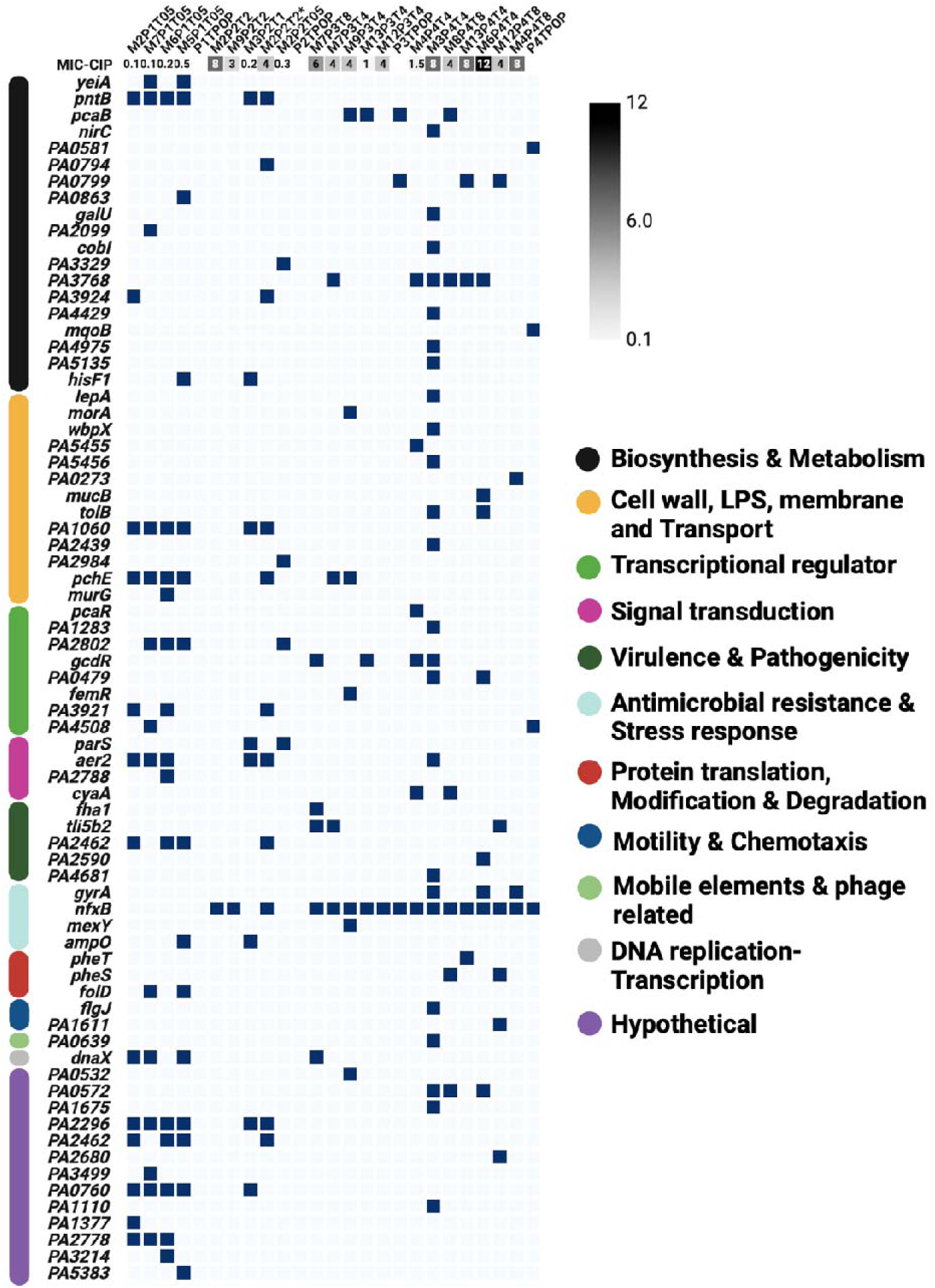
Genomic adaptation of *P. aeruginosa (*PAO1-*mCherry*-P*_CD_*-*gfp*+) to the murine lung in the presence of antibiotic pressure. Heatmap of mutational profiles in control isolates recovered across four sequential passages. Individual isolates are identified by mouse ID (M), passage number (P), and CIP-treated group designation (T), with corresponding ciprofloxacin (CIP) MIC values (mg/L) indicated above each column. Blue cells denote the presence of nonsynonymous mutations within specific genes, categorized by functional groups as indicated by the vertical color bars. * two isolates were collected from the same CIP plate but exhibited different MICs.

Consistent with previous findings (13), these *nfxB* mutants could be tracked in real-time via the emergence of GFP-positive colonies. The shift from large, mCherry-expressing ancestral colonies to smaller, stressed, slower- nfxB mutants expressing GFP- colonies correspond directly to the metabolic and growth trade-offs imposed by the development of high-level resistance (Fig. S5).

### Metabolic remodeling and genotype-phenotype association

The metabolic profiles from bacteria grown in LB medium revealed a progressive divergence between the control and treated evolved passages, culminating in a specialized metabolic state in the high-level resistant passage T4 (Fig. 5, Fig. S6). While control isolates maintained a relatively stable metabolic profile characterized by minor adaptations to the lung environment, the treated isolates exhibited a profound reorganization of central carbon and nitrogen metabolism starting as early as the second passage (T2) and reaching a specialized state by the final passage (T4). In the final passages (T3–T4), treated isolates exhibited a significant depletion of primary carbohydrates and energy precursors, including glucose, AMP, Trehalose, and Xylose (Fig. 5, deep blue). This shift correlates with mutations in transcriptional regulators such as *gcdR* (glucose catabolism) and the transhydrogenase *pntB* (redox balance), suggesting a redirection of carbon flux away from rapid growth toward maintenance and antibiotic defense. Notably, the high-level resistant isolates (T4) exhibited a significant accumulation of lipids and lipid-related metabolites in the exometabolome (Fig. 5, Fig. 6b). This increase in lipid signals is highly characteristic of *nfxB*-mediated resistance, reflecting the extensive membrane remodeling and metabolic costs associated with the constitutive overexpression of the MexCD-OprJ efflux system. The buildup of Formate and acetate in the later treated passages suggests a shift toward alternative or fermentative pathways, likely providing the biochemical basis for the reduced growth velocity and extended lag phase observed in our phenotypic assays (Fig. 2). The results represent a distinct metabolic canalization required to survive sustained ciprofloxacin stress.

**Fig. 5.**
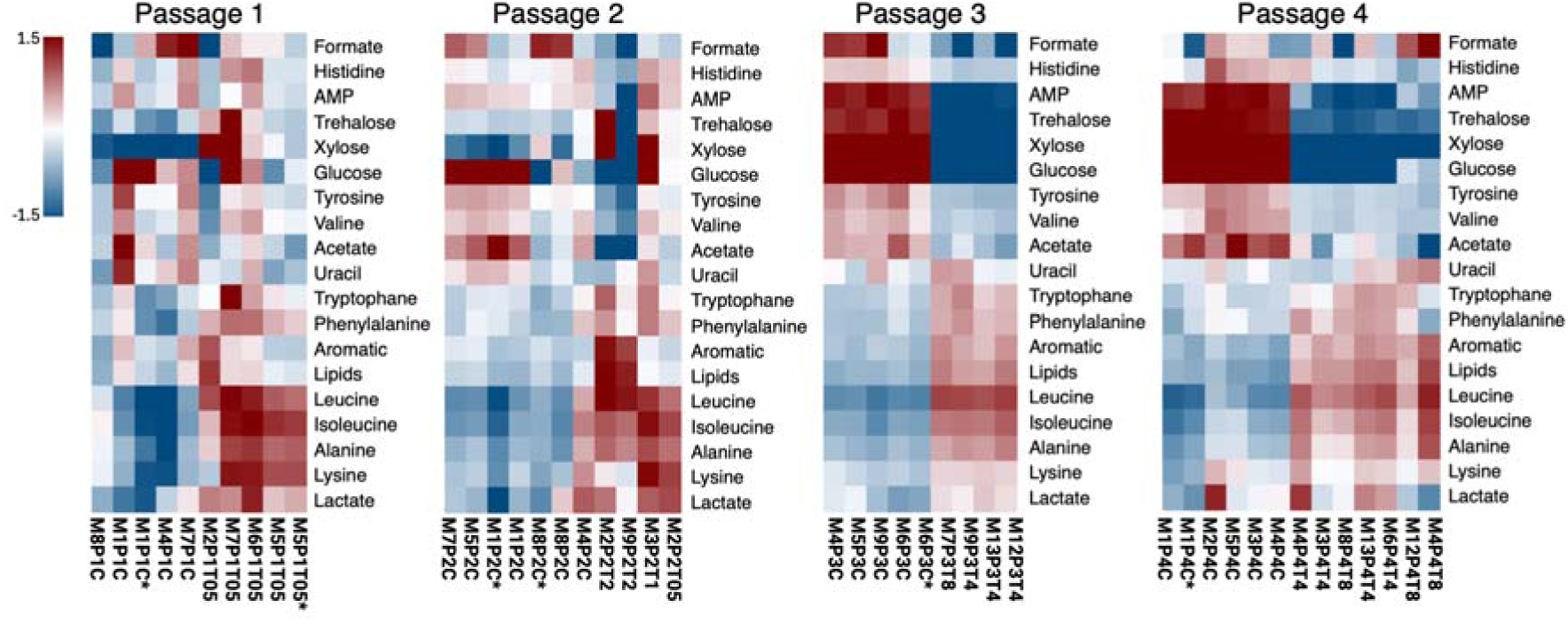
The metabolome of cell-free supernatants from *P. aeruginosa* strains from four adaptive evolution cycles was analyzed by ¹H-NMR spectroscopy. Cell-free supernatants were prepared from cultures grown in LB medium. Hierarchical clustering analysis of the NMR spectra clearly separated the samples into two distinct clusters between control strains and ciprofloxacin (CIP)-treated strains. The separation of two groups of metabolites was clear at P3 and P4. Metabolic analysis revealed that CIP-evolved *P. aeruginosa* strains at P3 and P4 exhibited higher sugar consumption (glucose, trehalose, and xylose) compared to control strains. The CIP-treated strains from P3 and P4 accumulated elevated levels of several amino acids, including tryptophan (Trp), phenylalanine (Phe), leucine (Leu), isoleucine (Ile), and alanine (Ala), as well as the glycolytic end product lactate (Lac). In contrast, lower levels of valine (Val), tyrosine (Tyr), and acetate (Ac) were detected.

**Fig. 6:**
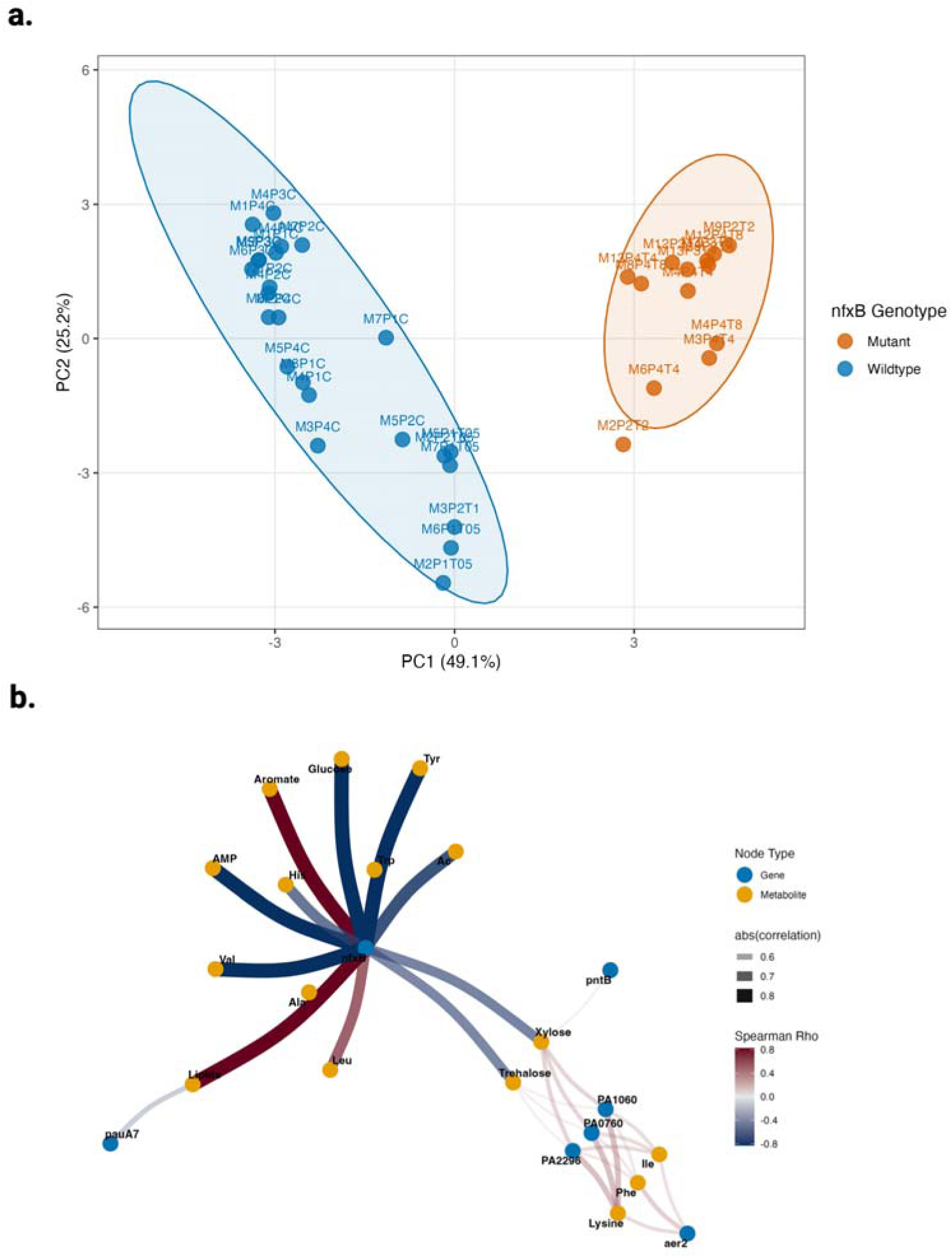
Integrated Genomic and Metabolomic Association Analysis. (a) Principal Component Analysis (PCA) of the endpoint exometabolome. Unsupervised PCA scores plot based on the H-NMR metabolic footprint of 37 *P. aeruginosa* isolates. The first principal component (PC1), explaining 49.1% of the total variance, reveals a profound metabolic divergence between high-level resistant lineages and control/early-passage isolates. Distinct clustering is supported by non-overlapping 95% confidence ellipses, with the primary separation driven by the emergence of the *nfxB* genotype in the treated groups. (b) Targeted Genomic-Metabolic Association Network. Visual representation of the non-linear dependencies between significant genomic variants and metabolite concentrations (nodes). The edge width represents the strength of the Spearman’s rank-order association (ρ), with the thickest lines denoting high-confidence associations (⎢ρ ⎢> 0.8). Red edges indicate a positive association (metabolite accumulation), while blue edges indicate a negative association (metabolite depletion). The network identifies *nfxB* as a central regulatory hub, showing strong negative associations with primary energy precursors (Glucose, AMP, Tyrosine). Secondary associations involving *pntB* and *pauA7* highlight the metabolic canalization required to support the physiological costs of sustained efflux activity, including significant associations with Lipids, Xylose, and Trehalose.

To statistically evaluate the relationship between emerging resistance genotypes and their corresponding metabolic landscape, we integrated the genomic and metabolic datasets into a unified association analysis (Fig. 6). Principal Component Analysis of the final endpoint metabolomic data confirmed a profound divergence in the metabolic states of the 37 isolates (used for both WGS and metabolic analysis) (Fig. 6a). PC1 (49.1%) clearly separated the isolates by *nfxB* genotype, supported by non-overlapping 95% confidence ellipses, while control and early-treated wild-type isolates clustered together. A targeted association network identified *nfxB* as a central hub, exhibiting negative associations with glucose, AMP, and tyrosine—the same metabolites observed to decline during the passage series (Fig. 6b, Fig. 5). Beyond resistance, the network links background adaptations to the metabolic profile. Mutations in *pntB* and *pauA7* show significant associations with xylose, trehalose, and lipids, suggesting these modifications facilitate the metabolic canalization required to support the high-energy demands of sustained efflux activity. An exhaustive association analysis revealed distinct functional modules (Fig. S7), and the full association matrix revealed distinct functional modules where specific genomic alterations consistently correlated with metabolite depletion or accumulation. A focused analysis of amino acid metabolism (Fig. S8) identified a high-confidence cluster involving *aer2, PA1060, PA2296,* and *PA0760*. These genes showed strong positive associations with lysine (ρ = 0.59), Isoleucine (ρ = 0.54), and Phenylalanine (ρ = 0.52), providing a mechanistic basis for the amino acid accumulation observed in the later passages of the study (Fig. S8, Fig. 5).

Further phenotypic analysis was done on the (T1-T4) isolates to determine whether this specialized metabolic and genomic state induced a significant shift in the broader antibiotic susceptibility profile, manifesting as collateral sensitivity to non-fluroquinolone antibiotics (Fig. 7). The MICs of CIP and the two antibiotics, tobramycin, an aminoglycoside, and aztreonam, a β-lactam, were determined and exhibited collateral sensitivity over passages. In passage 1, the isolates maintained baseline MIC levels as the ancestral strain (0.1 mg/L for CIP and 2 mg/L for tobramycin and aztreonam). While starting from the second passage, an inverse correlation emerged, as the isolates with higher CIP MICs showed lower MICs with tobramycin and aztreonam, even falling below the baseline MIC level (Fig.7). This indicates that the same evolutionary path driving CIP resistance simultaneously increased vulnerability to other drug classes.

**Fig. 7:**
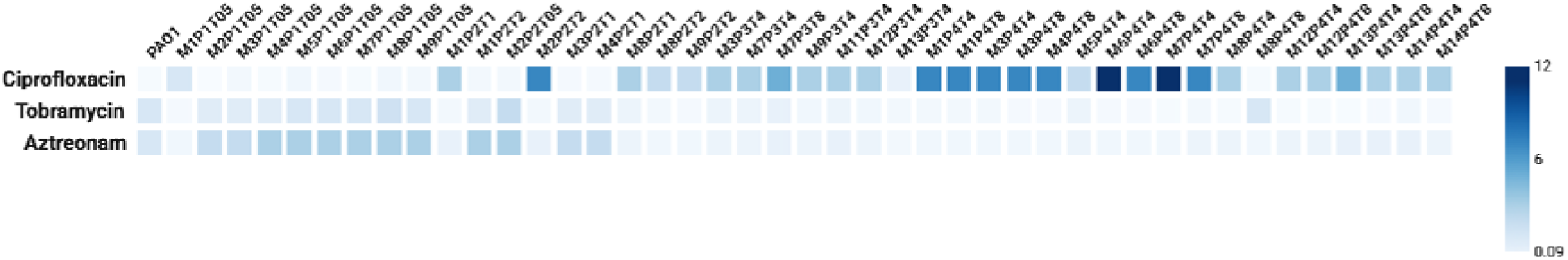
Collateral sensitivity. Heatmap showing the MICs (mg/L) for the three antibiotics, CIP, tobramycin, and aztreonam, that were determined using E-tests. Collateral sensitivity was observed over passages. The different numbers in the heatmap represent the different tested isolates. Individual isolates are identified by mouse ID (M), passage number (P), and CIP-treated group designation (T).

### Bacterial localization in the lungs and host response

Lungs from both untreated specimens and those subjected to CIP during the fourth passage were visualized using fluorescent in situ hybridization (FISH) to determine the existence of colonized bacteria on mouse lung sections (Fig. 8). The analysis revealed bacterial presence (in red; see arrows) in both control and treated groups. However, in the control lungs, bacteria were notably more concentrated and appeared condensed, primarily localized at the airway epithelium and spread across the tissue surface. Conversely, the treated lungs showed a reduced bacterial signal, possibly due to the CIP treatment. Additionally, some bacteria appeared to be intracellular, potentially engulfed by immune cells.

**Fig. 8:**
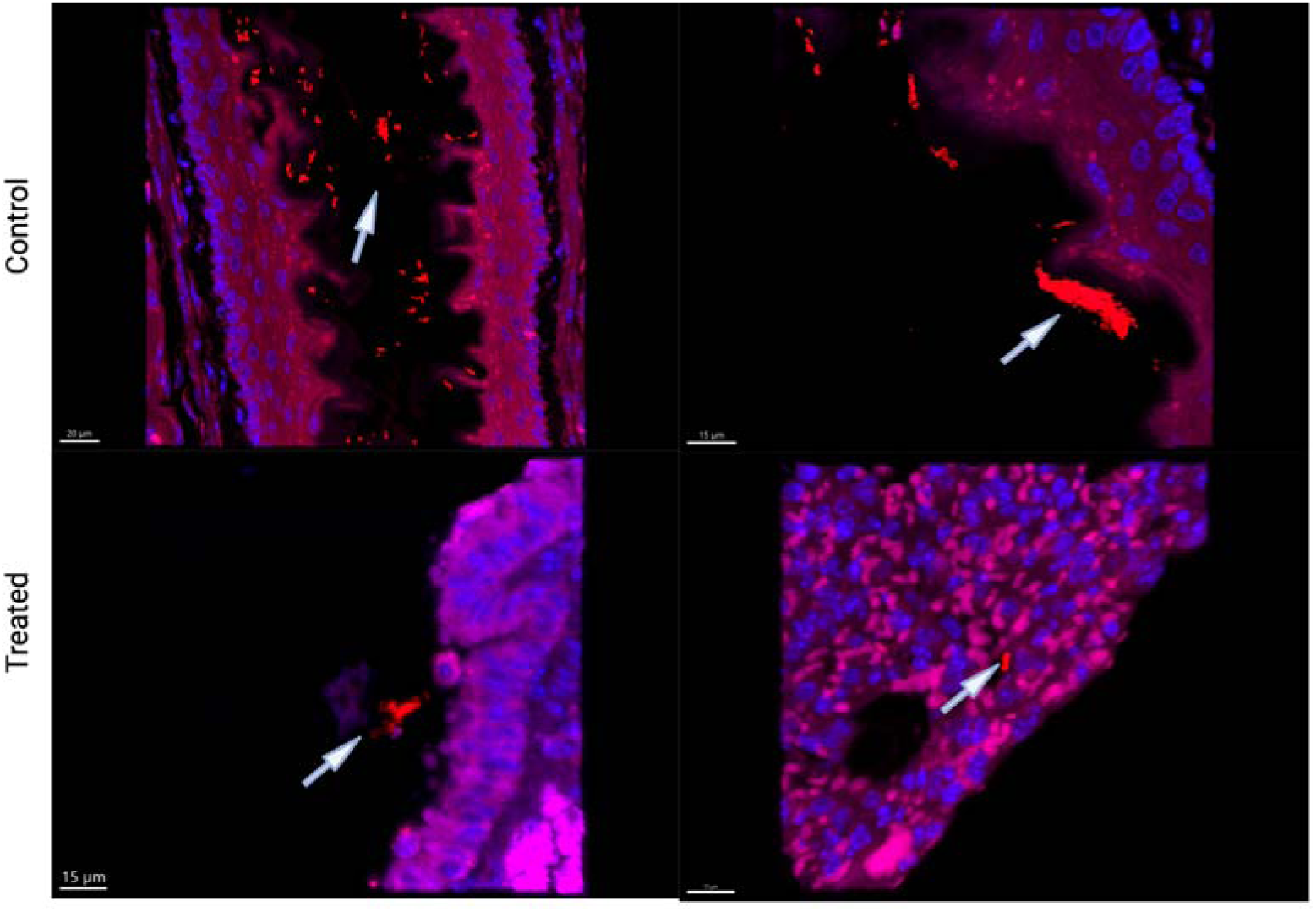
PNA-FISH staining of mice lung sections obtained from the fourth passage for both evolved groups, either CIP-treated or placebo (control). The images show bacterial cells are shown in red (TexasRed) conjugated probe and DAPI in blue as a nuclear stain. The images were captured by Zeiss LSM 880 confocal laser scanning.

The histopathological analysis of lung tissue sections revealed that chronic-like infection with *P. aeruginosa* embedded in alginate beads induced significant host tissue alteration compared to background controls (Fig.9 a, d). In the placebo-treated group control, we observed a high degree of neutrophilic inflammation characterized by dense leukocyte infiltration into the alveolar spaces and surrounding the bronchioles (Fig. 9b). Quantitative assessment confirmed a significantly higher inflammation score in placebo mice compared to those receiving CIP treatment (Fig. 9g; *p* = 0.011). Furthermore, the infection triggered significant differentiation of mucin-producing goblet cells within the bronchiolar epithelium, a hallmark of chronic airway irritation and mucus hypersecretion (Fig. 9e). CIP treatment significantly mitigated this response, resulting in a lower mucin score (Fig. 9h; *p* = 0.0496) and fewer visible goblet cells (Fig. 9f). These findings are highly consistent with our previous observations of smaller biofilm aggregates in CIP-treated mice, suggesting that while the bacteria evolved high-level resistance, the treatment-induced reduction in total biomass and altered metabolic state effectively lowered the overall inflammatory burden on the lung.

**Fig. 9:**
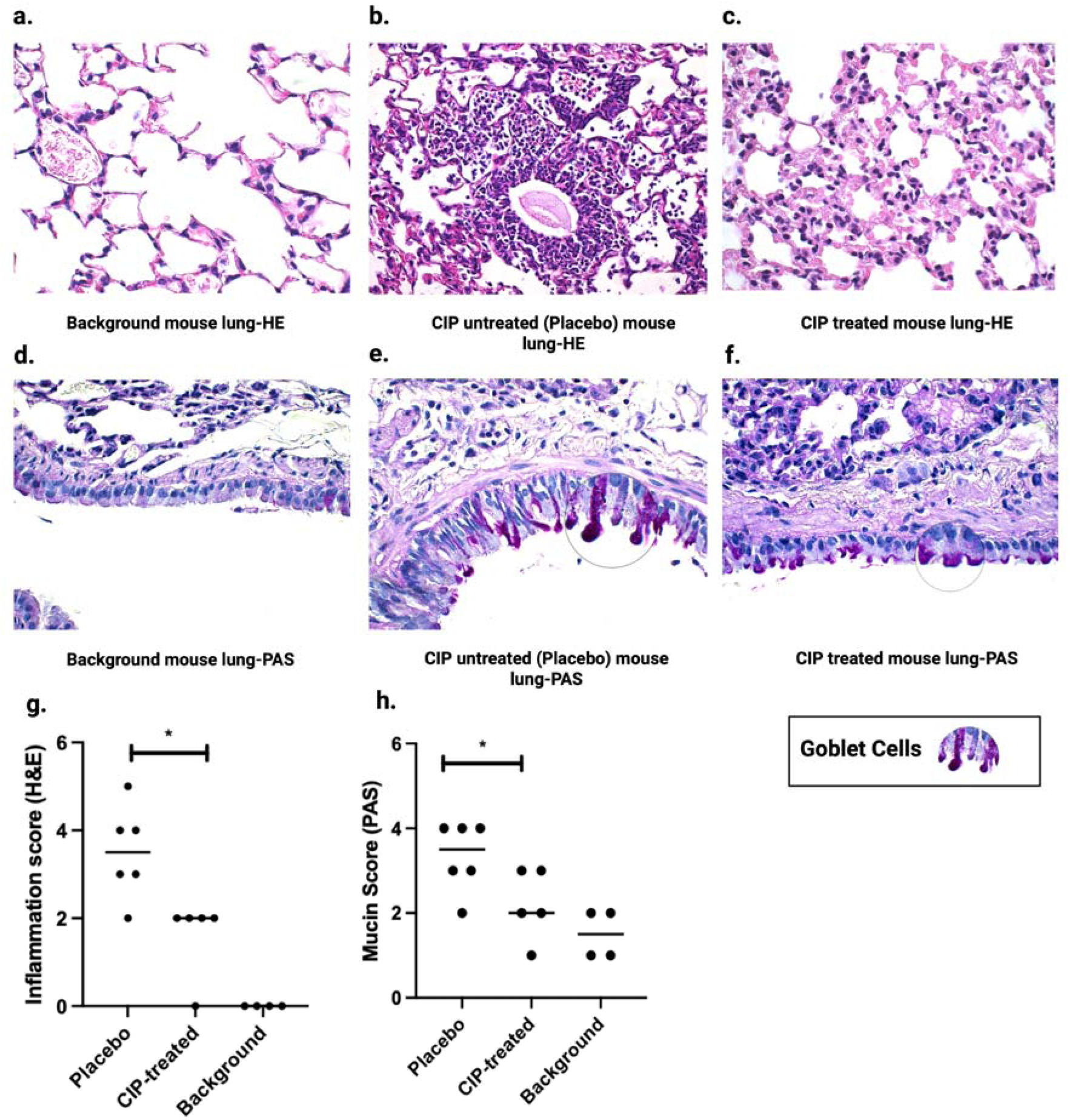
Histopathological analysis of host inflammatory response and mucus production. Representative lung sections from the fourth passage (T4) of evolution. (a–c) Hematoxylin and Eosin (H&E) staining of lung tissue from (a) Background (sterile alginate), (b) Placebo (untreated), and (c) CIP-treated mice. Untreated lungs (Placebo) exhibit high-grade neutrophilic inflammation and atelectasis, which are visibly attenuated in the CIP-treated group. (d–f) Periodic Acid-Schiff (PAS) staining highlighting mucus secretion. (e) Placebo lungs show significant goblet cell hyperplasia in the bronchioles (indicated by purple-stained mucin granules, circles), a feature reduced in (f) CIP-treated lungs. (g) Blinded histopathological inflammation scores based on H&E staining, demonstrating significantly lower tissue damage in CIP-treated mice compared to Placebo (*p* < 0.05). (h) Mucin scores based on PAS-positive goblet cell density, showing a reduction in mucus-producing cells following CIP-induced bacterial evolution (*p* < 0.05). All data represent individual mice from the final passage. Statistical significance was determined by an unpaired t-test (* *p* < 0.05). Scoring data per individual mouse lung is provided in Supplementary Table 1.

To further characterize the inflammatory environment within the lung, we performed a Luminex-based analysis of cytokine and protein concentrations in lung homogenate supernatants across all four passages. This analysis revealed an increased inflammatory response specifically in the treated mice during the first passage (T1), where levels of the neutrophil chemoattractant CXCL2 (Fig. 10a), the tissue remodeling protein MMP2 (Fig. 10b), and the pro-inflammatory cytokine IL-1β (Fig. 10d) were markedly elevated compared to both the first passage of the placebo-treated controls (*p* = 0.0025, 0.0464, and 0.02, respectively) and the background baseline. TNF-α (Fig. 10e) followed a similar pattern (*p* = 0.0037), which confirms a robust, treatment-induced early inflammatory burst that declines as the bacteria evolve resistance. This intense early signaling aligns with the high degree of neutrophilic infiltration observed in our initial histopathological sections. Interestingly, this inflammatory peak was not sustained; as the passages progressed and the bacteria transitioned toward a high-level, slow-growing resistance, the levels of CXCL2, MMP2, and IL-1β in the treated lineage significantly declined. By the third and fourth passages (T3–T4), these markers reached levels comparable to or even lower than the placebo-treated group, with slight divergence of IL-1β in the fourth passage, suggesting that the host’s innate immune response was no longer being aggressively triggered. This pattern was distinct from IFN-γ (Fig. 10f), which did not exhibit the same early treatment-specific elevation, indicating that the acute response was primarily driven by innate myeloid and structural signaling rather than a Th1-mediated adaptive response. The chemokine CCL2 (MCP-1) (Fig. 10g) showed a similar early elevated level for T1 passage (*p* = 0.0025), which decreased in successive CIP-treated passages, and inversely showed progressive increase in the later passages of the control group (P2C-P4C), suggesting a transition toward a more macrophage-heavy inflammatory infiltrate in the absence of antibiotic pressure. GM-CSF (Fig. 10h) showed a similar late-stage elevation specifically in the control isolates at passage four. This late-stage increase in CCL2 and GM-CSF in the placebo group suggests that untreated chronic infections continue to recruit and mature professional phagocytes, while the CIP-treated group exhibits a more quiescent immune profile in the final passages. The levels of G-CSF (Fig. 10i) (Granulocyte colony-stimulating factor) remained remarkably high across both treated and control groups throughout most passages, with numerous samples reaching the upper limit of detection (indicated by red dots). This indicates a massive and sustained systemic signal for granulocyte production, regardless of the treatment regimen.

**Fig. 10:**
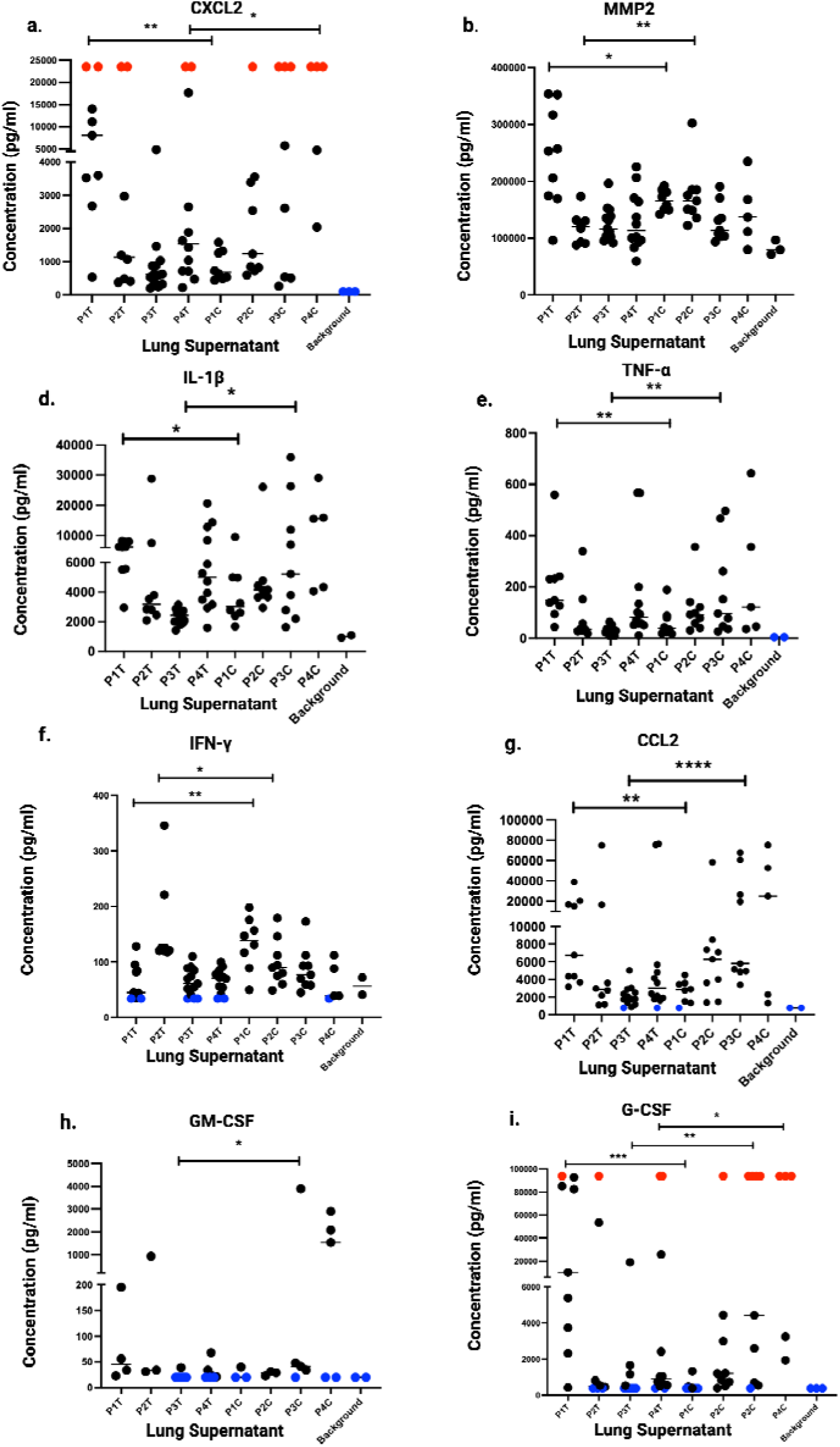
Inflammatory markers. a-i Cytokine levels in lung samples; concentrations (pg/ml) of cytokines, including (a) CXCL2, (b), MMP2, (c) IFN-γ (d) IL-1β (e) TNF-α, (f) G-CSF, (g) IL-1β, (h) CCL2, and (i) GM-CSF were measured in the supernatant of the evolved mice lung homogenates from CIP-treated (T) and control (placebo) mice (C) across four passages. Two-group comparisons were performed using the non-parametric Mann–Whitney test. Statistical significance is indicated as \**P*□<□.05, \*\**P*□<□.01, \*\*\**P*□<□.001, \*\*\*\**P*□<□.0001 using the non-parametric Mann–Whitney test. Values below the lower limit of quantification (LLOQ) are shown in blue, and values above the upper limit of quantification (ULOQ) are shown in red.

## Discussion

Understanding the evolutionary trajectories of bacteria within biofilms is critical for decoding the rise of clinical resistance and identifying exploitable therapeutic vulnerabilities. By implementing a prolonged, multi-passage CIP dosing regimen in a murine lung biofilm model, we successfully extended the selective window to capture the evolutionary continuum. This approach allowed us to trace the definitive transition from early, transient tolerance to stable, high-level resistance, circumventing the limitations of premature bacterial clearance that often hinder shorter-term *in vivo* studies.

This study shows that high-level resistance is not a solitary genetic event but a profound cellular transformation that imposes a significant fitness cost. We observed a distinct divergence in bacterial burden by the fourth passage, where CIP-treated populations exhibited lower CFU counts compared to control groups. This decline in population density coincided with a significant reduction in the maximum specific growth rate (*μ_max_*) and an extended lag phase in T4 isolates (Fig. 2b,c). In our previous study, the resistance threshold reached only 2 mg/L by the final passage (13); here, the intensified selective pressure accelerated this process, reaching 12 mg/L. These data suggest that the acquisition of high-level resistance forces a “metabolic bottleneck” where the bacteria trade kinetic growth for survival (21). Under the increased inoculum and in the absence of the antibiotic pressure, it could be that the bacteria gained adaptation to the lung environment, which led to increased persistence and growth over passages (22).

The evolutionary path under CIP selection was defined by a shift from early metabolic tolerance to the rapid development of high-level resistance. While isolates from the initial passages (T1–T2) showed mutations in diverse metabolic genes and increased survival at sub-MIC CIP concentrations compared to the ancestor strain, the second passage marked the critical emergence of *nfxB* mutations, which existed in all tested isolates by the fourth passage. WGS analysis revealed that these isolates reached a resistance threshold of 8–12 mg/L, driven by the accumulation of *nfxB* (regulating the MexCD-OprJ efflux pump) and primary target-site mutations in *gyrA*. While *nfxB* has been previously characterized as a metabolic modulator (23), our findings demonstrate the evolutionary canalization of this phenotype under sustained antibiotic stress within a complex host environment. We show that the metabolic cost of *nfxB*-mediated resistance is not a transient byproduct, but a fixed functional shift that dictates the success of the resistant lineage during the transition from tolerance to high-level resistance.

The trajectory of this adaptation follows a clear chronological pattern, while early adaptation in the treated lineage involved diverse mutations (e.g., *pchE*, *pntB*), the fixation of the *nfxB* mutation by the second passage triggered a profound “metabolic canalization” (23). Our integrated NMR and association network analyses identified *nfxB* as the primary driver of global metabolic variance, initiating a fundamental shift in carbon utilization. This was characterized by the systemic depletion of primary energy sources such as Glucose, Tyrosine, and AMP- reflecting a redirection of central carbon flux away from rapid growth and toward antibiotic defense.

This specialized state is further defined by two key signatures. First, our observation of lipid accumulation in the exometabolome (Fig. 6b) provides *in vivo* support for the hypothesis that long-chain fatty acids may serve as substrates or metabolic markers for the MexCD-OprJ efflux pump—a relationship previously suggested but not demonstrated in an evolutionary context (23). Second, the simultaneous accumulation of branched-chain amino acids, such as isoleucine and lysine, potentially driven by the *aer2/PA1060* cluster, indicates a specialized adaptation for survival in the nutrient-limited lung environment (23–25). Collectively, these results suggest that the *nfxB* metabotype is not merely a sign of “dysregulation” but a highly structured, evolved state required for persistence under host-imposed pressures.

Understanding how microbial resistance to one antibiotic affects susceptibility to other antibiotics is significant. Recent studies using lab evolution, genome sequencing, and functional analyses reveal that mutations causing multidrug resistance can also increase sensitivity to unrelated drugs by collateral sensitivity (26). This study provides a functional bridge between metabolic cost and clinical opportunity through collateral sensitivity. We observed that as passaged progressed and several isolates maintained the *nfxB* and *gyrA* mutations, they became sensitive to tobramycin and aztreonam. This shift represents a robust evolutionary trade-off (27). The mechanism likely stems from the physiological “Achilles’ heel” created by MexCD-OprJ overexpression (28). Previous research suggests that the metabolic and structural remodeling required for high-level ciprofloxacin resistance (such as changes in membrane potential, cell wall integrity, and down-regulation of other efflux pumps) facilitates the entry or efficacy of aminoglycosides and β-lactams (29, 30). By defining this “metabolic fingerprint” of resistance, we identify a specific window where the pathogen’s adaptation to one drug becomes its primary vulnerability to another, a concept with high translational potential for sequential therapy. Collateral sensitivity to tobramycin and aztreonam of CIP-resistant *P. aeruginosa* isolates has been shown to be a robust phenotype (31), raising the question of whether the collateral sensitivity effect can be leveraged to combat the drug resistance crisis and manage pathogens adapting to multiple antimicrobials. Resistant bacteria could potentially be controlled by switching to an alternative drug to which they exhibit collateral sensitivity (32).

Our host-response data reveal that the evolution of high-level resistance correlates with a reduction in host tissue damage. The CIP-treated group in our study is characterized by smaller biofilm aggregates and a reduced capacity to trigger host inflammation. This attenuation of virulence is a known consequence of *nfxB* mutations, which have been shown to impair the Type III Secretion System (T3SS) and increase susceptibility to host complement-mediated killing (23). The histopathological examination of the lungs showed PMN-dominated inflammation in untreated lungs and development of mucin-producing goblet cells, recapitulating the host-response to biofilm infection in CF patients (33).

The host’s immune profile mirrors this transition; the initial acute inflammatory increase (CXCL2, IL-1β, MMP2) in the treated group gave way to a quiescent state as the bacteria transitioned into their slow-growing, metabolic bottleneck (34). While the untreated control mice continued to drive chronic inflammation and recruit macrophages (evidenced by rising CCL2), the CIP-treated lineage evolved toward a less virulent, persistent phenotype, supporting the finding from our previous study (13). This suggests that in the lung, the price of resistance is paid in virulence: the bacteria survive the antibiotic, but they lose the kinetic velocity required to aggressively damage the host.

In conclusion, by integrating genomic, metabolic, and host-response data, we demonstrate that *P. aeruginosa* evolution is not a simple linear path, but a sophisticated adaptive strategy driven by the intense selective pressure within the lung microenvironment. The acquisition of high-level resistance via the *nfxB*-metabolic axis forces a transition into a specialized persistent state that is less aggressive to the host and uniquely vulnerable to collateral antibiotics. This view moves the field beyond simple MIC measurements and toward a deeper understanding of the “metabolic fingerprints” of resistance that can be exploited for personalized, translationally relevant therapeutic strategies.

## Acknowledgments

The authors thank Tina Wassermann and Louise Mørk for their technical assistance. D.H. acknowledges financial support from the Egyptian Ministry of Higher Education. This study was supported by a 2026 research grant from the European Society of Clinical Microbiology and Infectious Diseases (ESCMID) awarded to D.H.

## Author Contributions

**D.H.:** Writing – original draft, Visualization, Methodology, Investigation, Formal analysis, Data curation, Conceptualization. **K.C.W.**: Writing – review & editing, Visualization, Methodology, Investigation, Formal analysis. **L.B.:** Writing – review & editing, Methodology, Investigation. **S.S.P.:** Writing – review & editing, Visualization, Methodology, Investigation, Formal analysis. **P.R.J.:** Writing – review & editing, Visualization, Methodology, Investigation, Formal analysis. **C.M.:** Writing – review & editing, Supervision, Visualization, Methodology, Investigation, Formal analysis, Conceptualization. **O.C.:** Writing – review & editing, Supervision, Visualization, Methodology, Investigation, Formal analysis, Conceptualization. All authors provided final approval of the version to be published and agree to be accountable for all aspects of the work.

## Data availability

The datasets acquired from this study are available within the publication and the attached supplemental files. All the data for the sequencing results can be accessed through the SRA project nr: PRJNA1458751.

