## Supplementary data for "Evolutionary Trajectories of Ciprofloxacin Resistance in *P. aeruginosa* Lung Biofilms: Mutation Dynamics, Metabolomic Shifts, and Collateral Sensitivity"

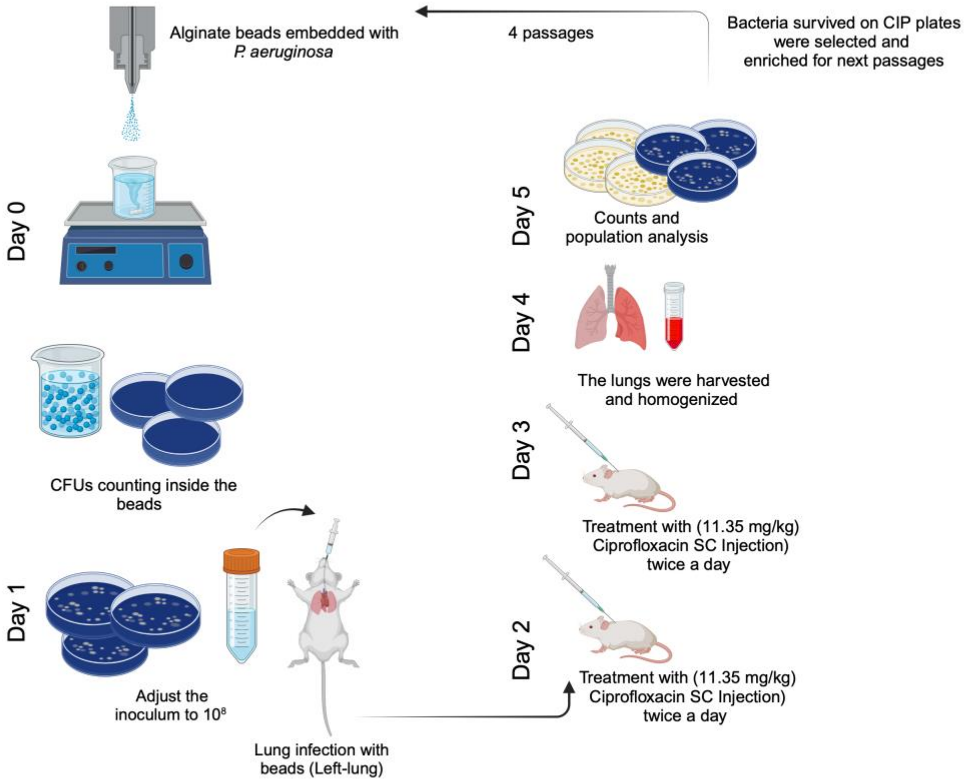


**S1** The experimental setup of the evolution of *P. aeruginosa* in a biofilm lung infection model in BALB/c mice; the experimental setup started with an overnight culture from a single colony of PAO1-*mCherry*-$P_{CD}$-*gfp*+; on Day 0, bacteria were embedded in alginate beads; on Day 1, mice were infected by injecting the left lung with alginate beads; on Day 2, the mice were treated twice with either CIP 0.25 mg (11.35 mg/kg) ×2 or saline (Placebo); on Day 3, the same treatment with CIP was repeated, and on Day 4 the mice were euthanized, and the lungs were collected, homogenized, and used for population analysis; on Day 5, colonies from CIP plates were selected for overnight cultures that were embedded in new alginate beads to infect a new group of mice (a new passage); the illustration was created with biorender.

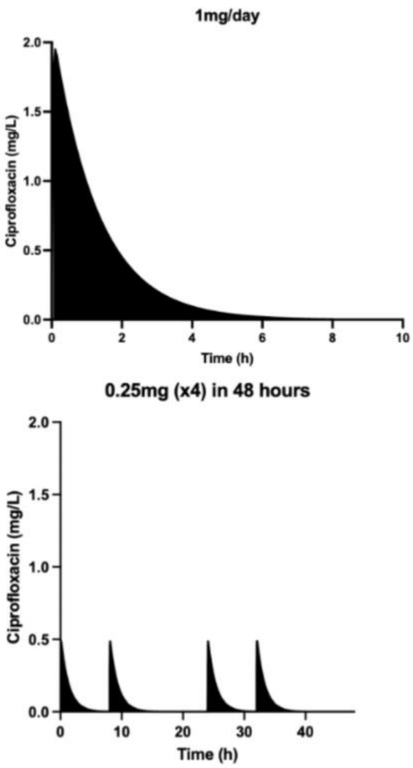


**S2.**Drug exposure and bacteriology. PK model fit for CIP administered at a dose of 1 mg (45.5 mg/kg) twice daily for 48 hours(1).

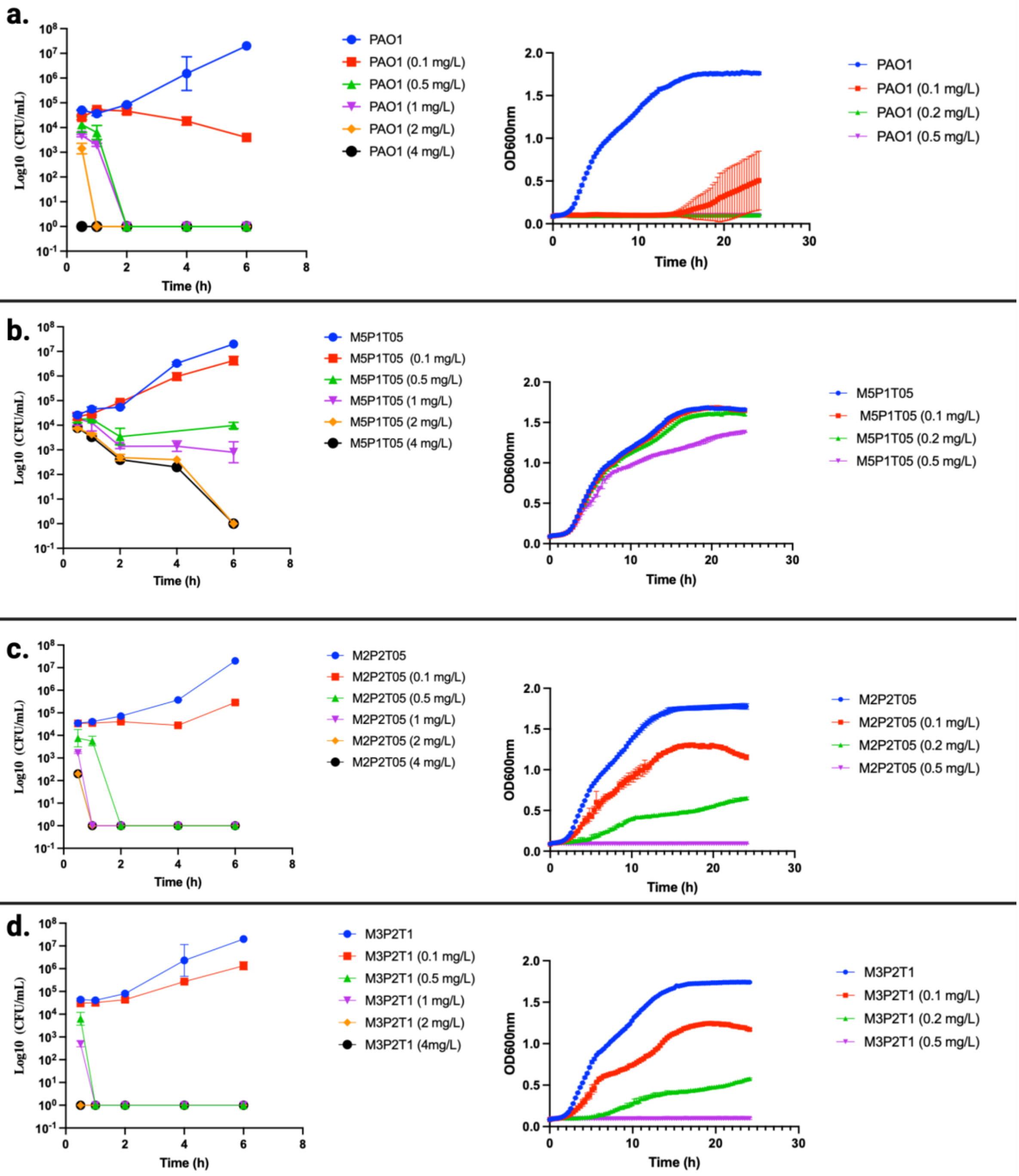


**Figure S3: a-d** Time-kill analysis (left) for the PAO1 ancestral strain and selected isolates from passages T1 and T2 at different ciprofloxacin (CIP) concentrations ranging from 0 to 4 mg/L. Corresponding 24-hour growth curves (right) for the same isolates under CIP concentrations ranging from 0 to 0.5 mg/L. Data represent mean SD from triplicate experiments.

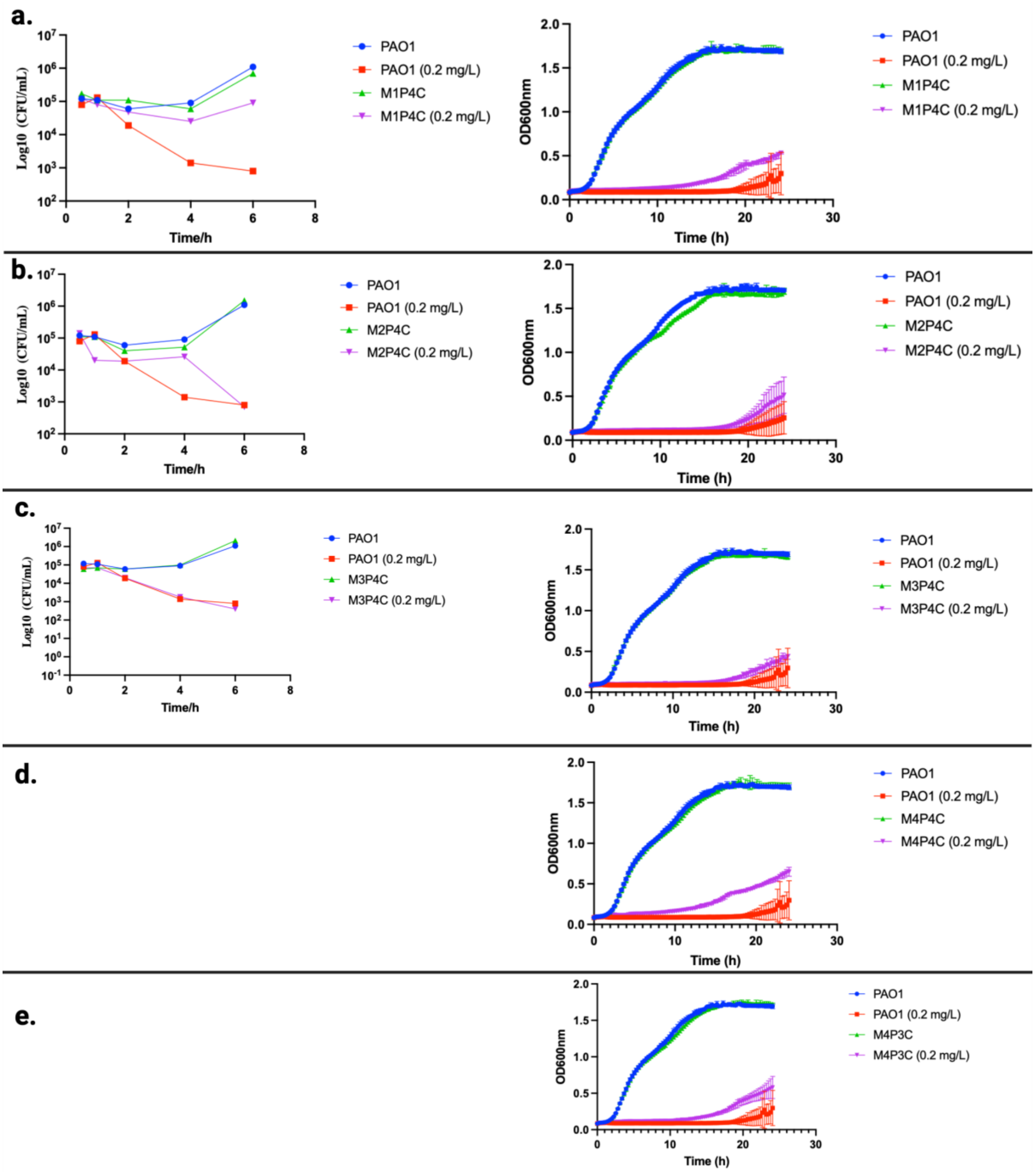


**Figure S4: a-d** Time-kill analysis (left) for the PAO1 ancestral strain and selected isolates from passages C1-C4 at CIP concentrations OF 0.2 mg/L. Corresponding 24-hour growth curves (right) for the same isolates under the same CIP concentration. Data represent mean  SD from triplicate experiments.

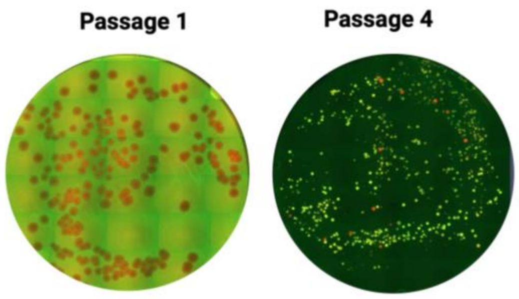


**S5**. The fluorescence observed in PAO1-*mCherry*-P$_{CD}$-*gfp*+ colonies from the lung bacterial populations of CIP-treated mice was observed over passages on LB agar plates free of CIP. Red fluorescence signifies the wild type, while green fluorescence represents *nfxB* mutants.

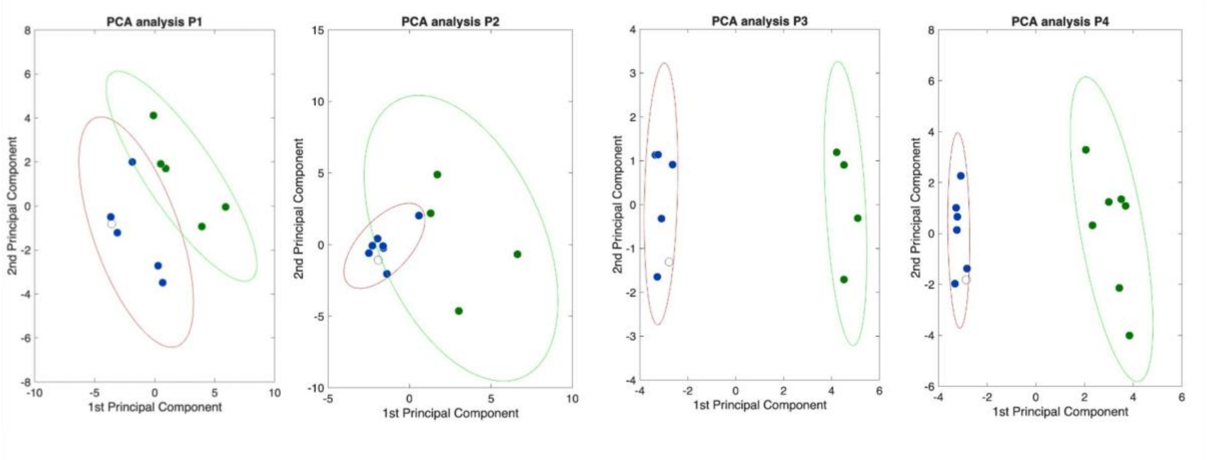


**S6.** PCA of the metabolites measured by $^1$H NMR for the different isolates from both Control and CIP-treated mice. Control (Blue), CIP-treated (green) and WT (open circle).

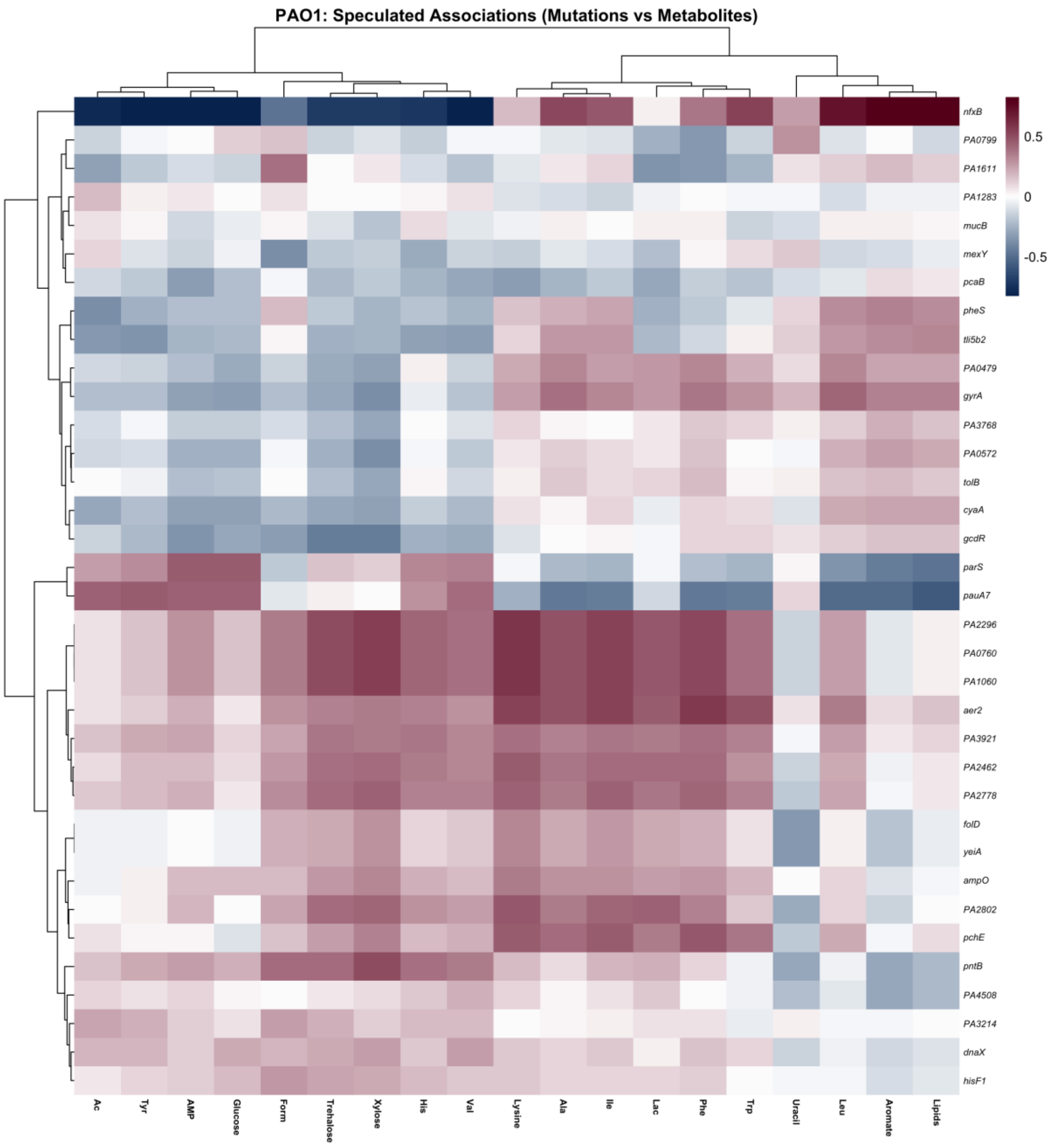


**S7: Full correlation landscape of genomic mutations vs. metabolites.** Comprehensive heatmap showing Spearman associations for all significant mutations across the tested isolates' metabolome.

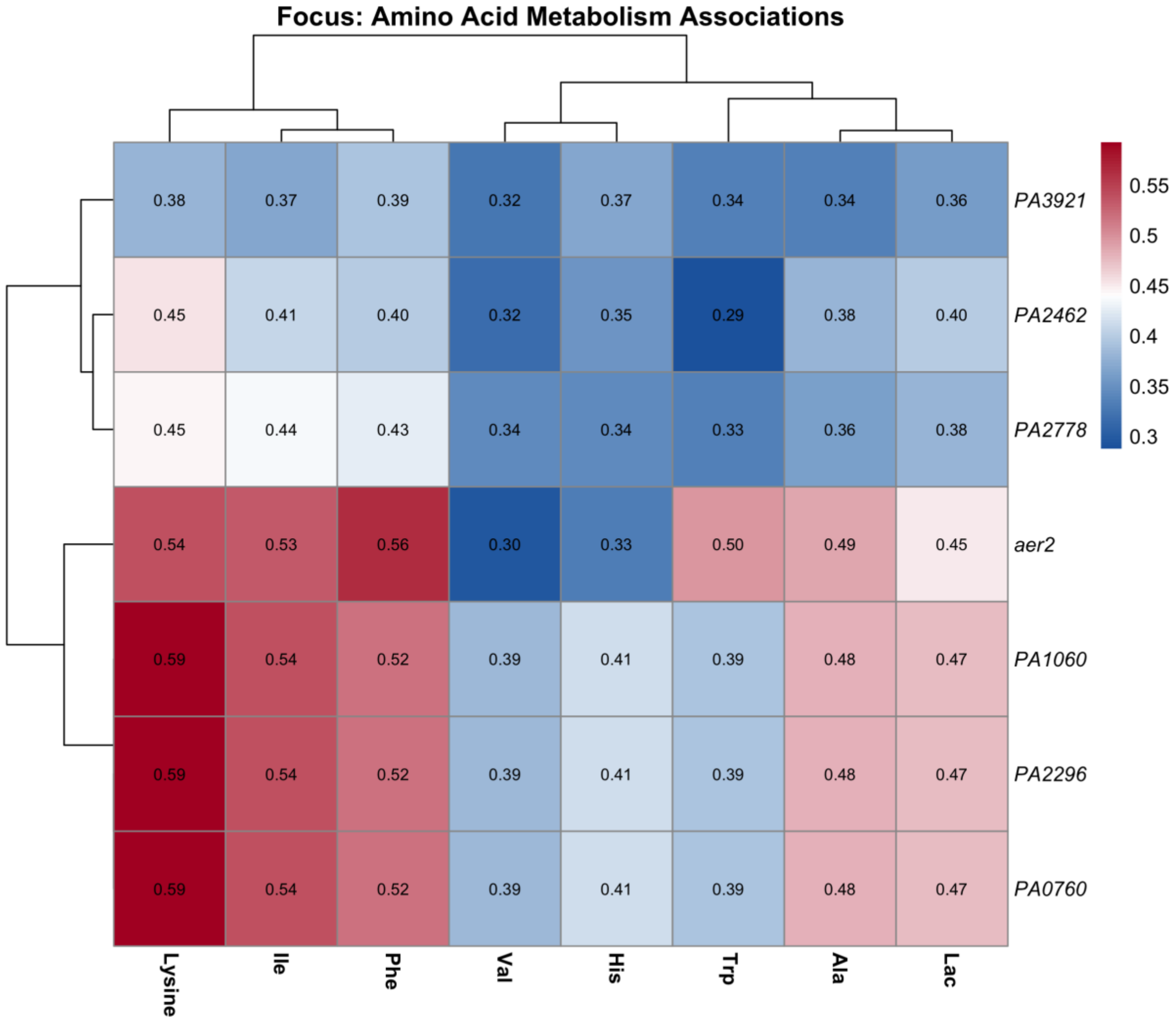


**S8: Focused correlation analysis of amino acid metabolism.** Targeted heatmap highlighting high-magnitude correlations between specific gene clusters (***aer2, PA1060***, etc.) and essential amino acids, with numerical values displayed.

| Mouse no. | Inflammation score | Mucin score |
|---|---|---|
| C1 | 4 | 3 |
| C2 | 3 | 4 |
| C3 | 2 | 2 |
| C4 | 4 | 4 |
| C5 | 3 | 3 |
| C6 | 5 | 4 |
| B1 | 0 | 1 |
| B2 | 0 | 1 |
| B3 | 0 | 2 |
| B4 | 0 | 2 |
| T1 | 2 | 3 |
| T2 | 2 | 1 |
| T3 | 2 | 2 |
| T4 | 2 | 2 |

Supplementary table1: Histopathological scoring of different mouse lungs. C= Control mice infected with *P. aeruginosa* embedded in alginate beads (Placebo), B=background mice inoculated with sterile alginate beads in their lungs that do not contain any bacteria, T= mice lungs exposed to CIP treatment.

1. Higazy D, Pham AD, van Hasselt C, Høiby N, Jelsbak L, Moser C, et al. In vivo evolution of antimicrobial resistance in a biofilm model of Pseudomonas aeruginosa lung infection. The ISME Journal. 2024;18(1).
